# Multi-Omic Atlas reveals cytotoxic phenotype and ROS-linked metabolic quiescence as key features of CTL-resistant HIV-infected CD4^+^ T-cells

**DOI:** 10.1101/2024.12.22.629960

**Authors:** Alberto Herrera, Louise Leyre, Jared Weiler, Noemi Luise Linden, Tan Thinh Huynh, Feng Wang, Colin Kovacs, Marina Caskey, Paul Zumbo, Maider Astorkia Amiama, Sandra Terry, Ya-Chi Ho, Doron Betel, R. Brad Jones

## Abstract

Cytotoxic T-lymphocytes (CTL) exert sustained pressure on reservoirs of HIV-infected cells that persist through years of antiretroviral therapy (ART). This selects for latently infected cells, but also potentially for cells that express HIV but possess intrinsic CTL resistance. We demonstrate that such resistance exists in HIV-infected CD4^+^ T-cells that survive rigorous CTL attack and map CTL susceptibility to cell identities and states defined by single-cell multi-omics and functional metabolic profiling. Cytotoxic CD4^+^ T-cells were prominently overrepresented amongst survivors, as were cells with quiescent metabolic profiles and low levels of reactive oxygen species (ROS) production. The induction of ROS production by treatment with deferoxamine sensitized these cells to CTL-mediated elimination. Reservoir-harboring cells from clinical samples share the above transcriptional features, being enriched for quiescent states. Our results provide an atlas for elucidating features of CTL resistance in HIV reservoirs, and identify oxidative stress as a therapeutic target to facilitate reservoir elimination.

## INTRODUCTION

HIV, although manageable as a chronic condition, currently lacks a safe or scalable cure. Antiretroviral therapy (ART) effectively suppresses HIV replication, preventing disease progression and transmission to sexual partners. However, ART does not affect established ’reservoirs’, which are rare cell populations harboring integrated, intact proviruses that can reignite viral replication if ART is discontinued^1–3^. A critical reservoir exists in memory CD4**^+^** T-cells, which become increasingly clonal over time due to infected cell proliferation^4–6^. These clonal dynamics facilitate Darwinian selection, favoring reservoir-harboring cells that exhibit fitness advantages in the evasion of immune effectors, including HIV-specific cytotoxic T-lymphocytes (CTL)^7–12^.

One key mechanism enabling these reservoir-harboring cells to evade CTL clearance is viral latency^1,13^. Amongst intact proviruses, those harbored in genomic locations or orientations that hinder viral transcription increasingly dominate over time on ART^10,14^. Latency is not complete, however. Cells expressing low to moderate levels of HIV RNA can be detected after decades of treatment, as can low levels of HIV p24 protein in blood plasma^15,16^. Associations between virologic measures of HIV persistence and HIV-specific T-cells frequencies and functions suggest ongoing CTL recognition of infected cells^17–21^.

Two main mechanisms likely contribute to the persistence of HIV-expressing cells despite ongoing CTL surveillance. First, clonal expansion of HIV-infected CD4**^+^**T-cells on ART replenishes fractions of the population that express antigen and are cleared^22–24^. Second, some infected CD4**^+^** T-cells may be intrinsically resistant to CTL attack, mirroring a phenomenon observed in cancer (reviewed in ^25^). Overexpression of the anti-apoptotic factor BCL-2 has been identified as one mechanism of CTL resistance^26–28^. The BCL-2 antagonist ABT-199 sensitizes HIV reservoir-harboring cells to CTL elimination *ex vivo* and is currently being investigated in a clinical trial^26^ (NCT05668026). Multi-omic profiling of *ex vivo* HIV reservoir-harboring cells has identified other potential mechanisms of CTL resistance, including the granzyme B inhibitor SERPINB9, and the immune checkpoint ligand PVR (CD155)^12,22,29^. However, understanding the contribution of these and other factors to HIV persistence has been limited by the lack of a reference profile of CTL-resistant, HIV-infected CD4**^+^**T-cells.

In this study, we confirm the existence of HIV-expressing primary CD4**^+^** T-cells that can survive repeated CTL attacks. We then characterize their transcriptomic and surface protein profiles at the single-cell level, providing a valuable reference for reservoir characterization studies. Furthermore, we uncover how metabolic and oxidative stress features influence the susceptibilities of these cells to CTL attack and show that an FDA-approved drug can be leveraged to boost the elimination of HIV-infected cells *in vitro*. This work shifts the paradigm on HIV persistence and opens novel therapeutic avenues for curing infection.

## RESULTS

### A rare population of HIV-expressing memory CD4^+^ T-cells resists repeated CTL attacks

CD4**^+^** T-cells are highly diverse with respect to both their maturational (e.g. naive versus memory) and polarization (e.g. Th_1_ versus Th_17_) profiles. While it would be of interest to study how intrinsic susceptibility to CTL varies as a function of these parameters, we reasoned that mechanistic drivers of resistance were more likely to be revealed by their fluctuations and selection within a relatively homogenous starting population. We therefore adapted a method of activating and culturing naïve CD4**^+^** T-cells under non-polarizing conditions to foster a central memory T-cell (T_CM_) phenotype, which we infect with HIV during the activation process (Figures 1A, S1A and S1C-D)^30,31^. To enable rigorous evaluation of specific CTL killing, we developed a 2- virus system where cells are infected with either a wild-type (WT) clinical isolate strain of HIV, ’HIV_JRSCF_’ (targets)^32^, or a version of this same virus with a single escape mutation in the TW10- Gag epitope, HIV_JRSCF TW10esc_ (bystanders), preventing recognition of the epitope by TW10-specific CTL clones generated from the peripheral blood of people living with HIV (PLWH) (Figures S1B and S1E).

**Figure 1.**
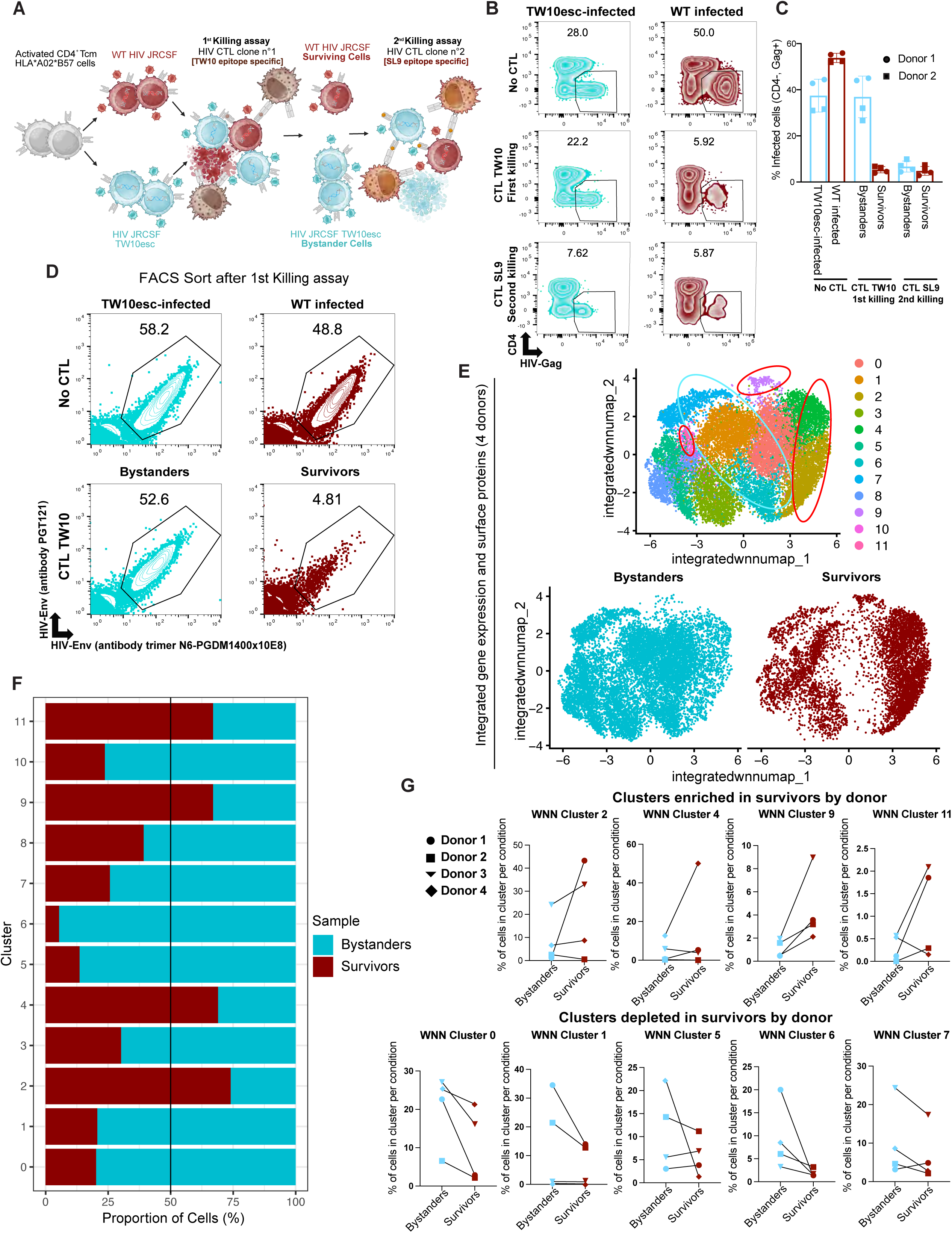
Existence and multi-omics profiles of CTL-resistant HIV-expressing CD4^+^ T-cells. **(A).** Schematic of ’sequential killing assay’, featuring two rounds of killing with CTL clones targeting the Gag-TW10 or Gag-SL9 epitopes. **(B and C**). Representative (**B**) and summary (**C**) flow cytometry data showing frequencies of HIV-infected cells (CD4^-^, HIV-Gag**^+^**) cells from different stages of sequential killing assays. In first-round killing with a Gag-TW10-specific CTL clone the majority of cells infected with WT HIV_JRCSF_ (red) are killed, whereas those infected with escape mutant HIV_JRSCF TW10esc_ (blue) are not. In second-round killing, ’bystanders’ from the first round (HIV_JRSCF TW10esc_) are now efficiently killed whereas ’survivors’ HIV_JRCSF_ resist further elimination. Shown in **C** are fold changes in infected (CD4^-^, HIV-Gag^+^) cells pre- and post-killing across two donors in technical duplicates. **(D)** Gates used to sort infected cells from bystander and survivor populations for analysis by ECCITE-seq. Populations containing WT HIV_JRCSF_ or HIV_JRSCF TW10esc_ were first gated based on CFSE and CTV labeling, respectively (Figure S1F-G). Within each of these, cells actively expressing HIV were identified and sorted based on co- staining with the HIV-Env specific broadly neutralizing antibodies PGT121 and N6- PGDM1400x10E8. **(E)** Harmonized WNN UMAP projections of bystander and survivor cells after CTL exposure from 4 donors clustered using cell-to-cell multimodal neighbors (weighted combination of RNA and surface protein similarities (n = 17,231 cells). Donors 1, 2 and 4 were HIV-negative individuals while donor 3 was a PLWH. **(F**) Proportion of bystander and survivor cells across WNN clusters. >50% indicates preferential resistance to CTL whereas <50% indicates preferential susceptibility. **(G)** Proportion of bystander or survivor cells belonging to each WNN cluster out of total bystander or survivor cells stratified by individual donor.

We first assessed whether a CTL resistant population exists within these infected T_CM_, using a ’sequential killing assay’. Infected T_CM_ were generated using PBMCs from two HLA- B57^+^A02^+^ people without HIV. Cells infected with either WT HIV_JRSCF_ (targets) or HIV_JRSCF TW10esc_ (bystanders) were stained with different dyes, mixed together and co-cultured with an HLA-B57- restricted CTL clone targeting the Gag ’TSTLQEQIGW’ (TW10) epitope for 16 hours (Figures 1A and S1D-S1E). This design ensures that both infected populations experience the same milieu, which is important since factors such as IFN-γ can alter susceptibility to CTL killing (e.g. by upregulating MHC-I)^33^. As expected, HIV_JRSCF TW10esc_ -infected cells (bystanders) were only minimally affected by the presence of the TW10-specific CTL clone, while WT HIV_JRSCF_ -infected cells (targets) were mostly eliminated (approximately 90% of initial HIV-Gag**^+^** cells) (Figures 1B and 1C). A small population of targeted cells, however, survived this initial CTL attack (10% of initial HIV-Gag**^+^** cells) (Figures 1B and 1C). To determine whether this rare population of surviving targets was intrinsically resistant to CTL killing, we performed a second round of CTL co-culture, where now WT HIV_JRSCF_-infected survivors and HIV_JRSCF TW10esc_-infected bystanders were both subject to attack. This was achieved by removing the TW10-specific CTLs from the first round of killing and co-culturing the infected cells with an HLA-A02-restricted, SL9-Gag-specific CTL clone (Figure 1A). The SL9-specific CTLs efficiently eliminated the HIV_JRSCF TW10esc_-infected cells that had been bystanders in the first round of killing, indicating efficient cytotoxic effector function. Critically, however, the WT HIV_JRSCF_-infected cells that had been targeted and survived in the first round, were not further reduced by the SL9-specific CTL attack (Figures 1B and 1C). This provides direct evidence for the presence of rare populations of HIV antigen-expressing CD4**^+^** T- cells that are intrinsically resistant to CTL-mediated killing.

### Distinct clusters of CTL-resistant cells identified by multi-omic profiling

Having confirmed the ability of this *in vitro* system to identify *bona fide* CTL resistant cells, we next characterized the transcriptional profiles, the surface expression of 140 immunologic markers, and the clonotypic identities of rare CTL-resistant survivors compared to bystanders using Extended CRISPR Compatible Cellular Indexing of Transcriptomes and Epitopes by Sequencing (ECCITE-seq)^34,35^ (Figures 1D and 1E). For both populations, we sorted cells based on HIV-Env expression to isolate cells that survived despite antigen expression, versus selecting for latently infected cells (Figure 1D). Harmony-based integration of ECCITE-seq data from 4 donors, followed by Weighted Nearest Neighbor (WNN) analysis of integrated gene expression and surface protein profiles revealed 11 different unsupervised clusters among total HIV-expressing T-cells. Strikingly, CTL-resistant survivors were specifically enriched in 4 of the 11 clusters; clusters 2, 4, 9 and 11 (Figure 1F and Table S1). In breaking down the skewing of survivors and bystanders in these clusters by donor, we observed that 3/4, 2/4, 4/4 and 3/4 donors showed enrichment of survivors versus bystanders in clusters 2 (average 2.5-fold enrichment), 4 (average 3-fold enrichment), 9 (average 3.9-fold enrichment) and 11 (average 3.6-fold enrichment) respectively (Figure 1G and Table S1).

Conversely, certain clusters of infected cells were highly sensitive to elimination (skewed towards bystanders). These were clusters: 0 (1.9-fold reduction), 1 (2.0 fold-reduction), 5 (1.9- fold reduction), 6 (5.3-fold reduction), and 7 (1.5-fold reduction) with skewing observed in 4/4, 2/4, 2/4, 4/4 and 3/4 donors respectively (Figure 1G and Table S1). While we also leveraged TCR sequencing to consider whether particular T-cell clones varied in their susceptibilities to killing, we observed very little in the way of expanded clones in either the bystander or survivor populations (Figure S2A). This is unsurprising given that our model uses polyclonal activation of naïve CD4 T-cells, with limited time for clonal expansion.

### An atlas of transcriptional features defining CTL-resistant or susceptible clusters

To probe for transcriptional programs that may contribute to CTL resistance or sensitivity in the cells making up survivor- and bystander-enriched clusters, we identified the top differentially expressed genes in each cluster and performed pathway enrichment analysis. Amongst the CTL-resistant clusters, we identified a cytotoxic signature and high expression of adhesion genes in cluster 9, high expression of ribosomal proteins in cluster 2, and an interferon stimulated gene (ISG) signature/high antiviral response in cluster 4 (Figure 2A and 2B). Conversely, amongst CTL- sensitive clusters, we identified high expression of activation and stress genes in cluster 0; activation and metabolic genes in cluster 1; activation, proliferation and adhesion genes in clusters 5 and 6, and cell division genes in cluster 7. Clusters 3 and 10 were omitted from further analysis due to their cells expressing a disproportionately high percentage of mitochondrial genes compared to other clusters, indicating these were lower quality/pre-apoptotic cells (Figure S2B).

**Figure 2.**
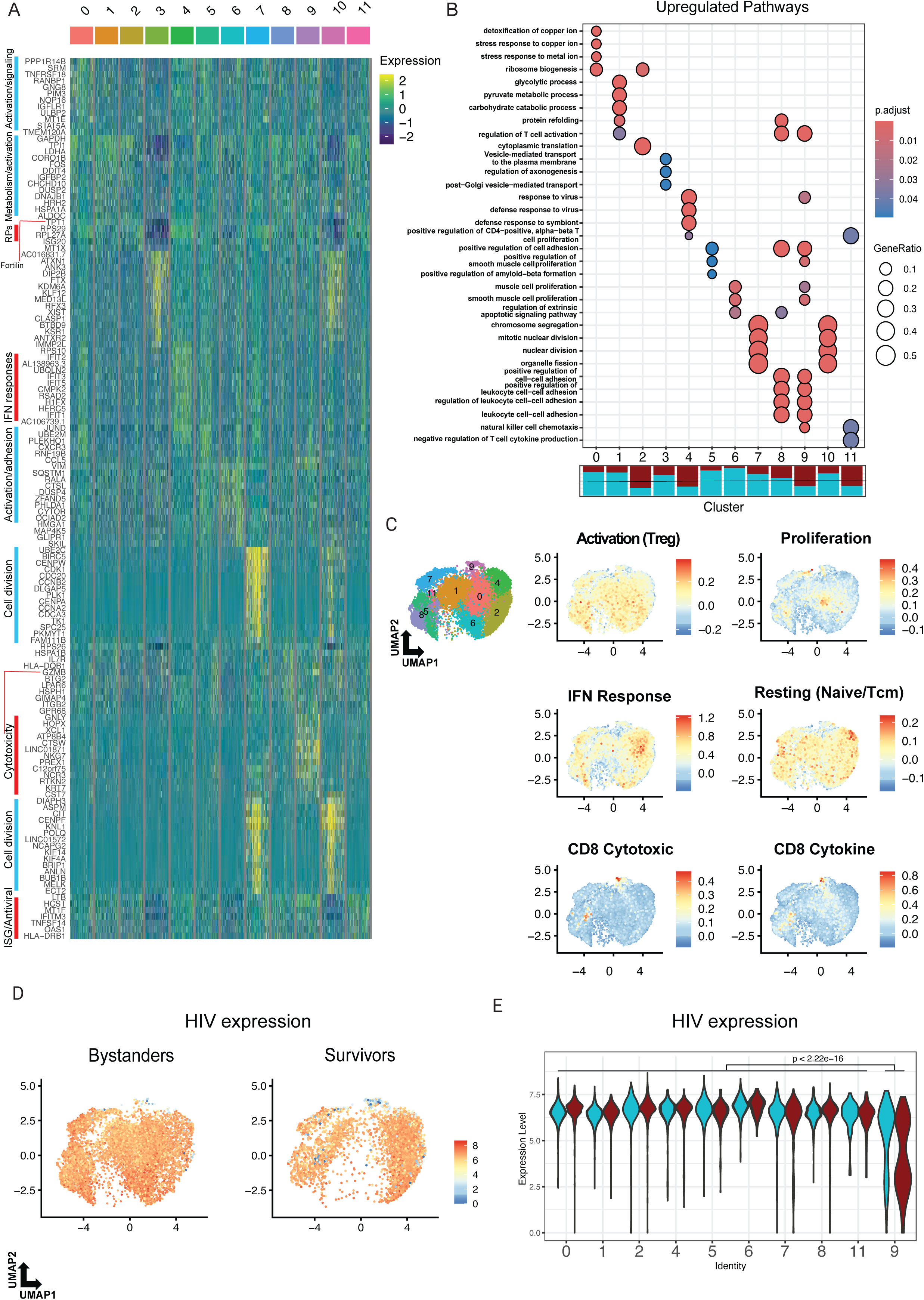
Single cell atlas of transcriptional features defining CTL-sensitive and CTL- resistant states in infected cells. **(A)** Heatmap of top 10 differentially expressed genes with a log_2_ fold-change difference of at least 0.25 passing min.pct ≥ 0.25 and P_adj_ < 0.05 among all WNN clusters highlighting overexpressed genes from CTL sensitive clusters (blue) and CTL-resistant clusters (red). **(B)** Top 5 gene ontology enriched pathways with a false discovery rate of 0.05 using gene ontology BP gene-sets using overexpressed genes by each WNN cluster passing the filters described in **(A)**. **(C)** Module scoring of T-cell activation and functional states defined by Szabo *et* al., 2019^36^, mapped to WNN clusters. Lower panels indicate that cluster 9 has a cytotoxic profile. **(D)-(E)** Quantification and comparison of HIV transcript expression among clusters. P values of Wilcoxon tests are shown.

To further define the identifies of bystander- and survivor-enriched clusters, we leveraged an independent dataset defining activation status and functionality modules in T-cells and overlaid these on our WNN-defined clusters^36^. We scored the levels of gene expression from infected cells on modules from this reference dataset descriptive of activation profiles: “Treg-like activation” and “Proliferation”; an intermediate activation module: “IFN response”; cytotoxicity profiles: “CD8 cytotoxicity” and “CD8 cytokines”; and a resting “naïve/T_CM_ CD4**^+^** T-cells” profiles. In line with our previous analysis, CTL-sensitive clusters 0, 1, 5, 6 and 7 had the highest expression of modules highly associated with activation, such as the “Treg-like activation” module in clusters 0, 1, 5 and 6, and the “proliferation” module in clusters 0 and 7 (Figures 2C and S2C-S2D). For CTL-resistant clusters, cluster 4 had high expression of “resting naïve/T_CM_ CD4**^+^** T-cells” as well as the “IFN response” modules, while cluster 9 scored markedly high for the “CD8 cytotoxicity” and "CD8 cytokine” modules (Figures 2C and S2E-S2H). These results demonstrated a tendency for cells with higher activation and proliferation profiles to readily succumb to killing by CTL, while cells enriched for resting, IFN response and cytotoxic profiles tended to be resistant to CTL.

In addition to host genes, we assessed levels of HIV transcription in relation to the CTL susceptibilities of cell clusters. Since cells had been sorted for those expressing HIV-Env, essentially all cells contained readily detectable levels of HIV transcripts. While CTL-resistant clusters 2 and 4 neither showed lower basal levels of HIV transcripts than susceptible clusters in the unselected bystanders, nor selection for the lower end of this distribution amongst survivors, cytotoxic cluster 9 stood out in having significantly lower levels of HIV transcripts than other clusters - manifesting as a bi-modal distribution of HIV-transcript-high versus HIV-transcript-low, and the latter was moderately enriched amongst the survivors of CTL selection (Figures 2D-2E). Thus, overall, levels of HIV expression did not appear to be a major determinant of CTL susceptibility (in the context of our model of sorted HIV-Env**^+^** cells). However, relatively low levels of HIV expression might contribute to CTL evasion in a subpopulation of cytotoxic HIV-infected CD4**^+^** T-cells.

Our results indicate that amongst infected cells generated in a cultured T_CM_ system we can capture diversity in cellular and immunophenotypic modules associated with both resistance and susceptibility to CTLs. The strongest transcriptional predictor of CTL-susceptibility in HIV-expressing CD4**^+^** T-cells appears to be high expression of activation and proliferation programs, while CTL-resistant cells were enriched for multiple cellular states including IFN responses and engagement of a cytotoxicity program.

### CTL resistance in HIV-expressing cells is linked to quiescent transcriptional and immunophenotypic states

To pan-out to the most widely shared cellular features delineating CTL-resistant survivor cells from bystanders, we next performed differential gene expression and pathway enrichment analysis on all survivor cells compared against all bystander cells, across the 4 donors. Amongst the genes with the most reduced expression levels in survivors were those coding for TNF-receptor superfamily member costimulatory receptors *TNFRSF18* (GITR), *TNFRSF4* (OX40) and *TNFRSF9* (4-1BB), the chemokine receptor *CXCR3*, the dual specificity phosphatase *DUSP4* and the metallothionine *MT2A* – all transcripts expressed downstream of T-cell activation^37–39^ (Figure 3A and Table S2). Pathway enrichment analysis revealed that several aerobic metabolism-related gene-sets were the most downregulated in survivor cells (Figure 3B). Relatedly, oxidative stress (KEAP1-NERF2) as well as proliferation (cell cycle) gene-sets were robustly down in these cells compared to bystanders. As T-cell activation induces a shift towards aerobic glycolysis and heavy reliance on glucose oxidation to sustain clonal expansion, termed the Warburg effect^40^, we proceeded to quantify the extent to which genes along the glucose metabolism cascade were differentially expressed. We calculated module scores in single cells based on their individual expression levels of genes annotated as members of each pathway subtracted by aggregate control features^41^. Our analysis revealed significant downregulation in survivor cells of glycolysis, aerobic respiration, and oxidative phosphorylation pathways as well as other general metabolism sets (Figure 3C and S3A-S3D). Oxidative stress, arising from the production of reactive oxygen species (ROS) as a natural consequence of active metabolism^42,43^, was found to be reduced in survivors based on our pathway enrichment analysis, which revealed downregulation of genes involved in the oxidative stress response (Figure 3B). Accordingly, module scoring of ROS and and oxidative stress response pathways revealed concomitant downregulation of their gene-sets in survivor cells (Figure 3C), indicating that lower activation thresholds in survivors translated to a more quiescent-like transcriptional program in these cells with reduced levels of oxidative stress responses.

**Figure 3.**
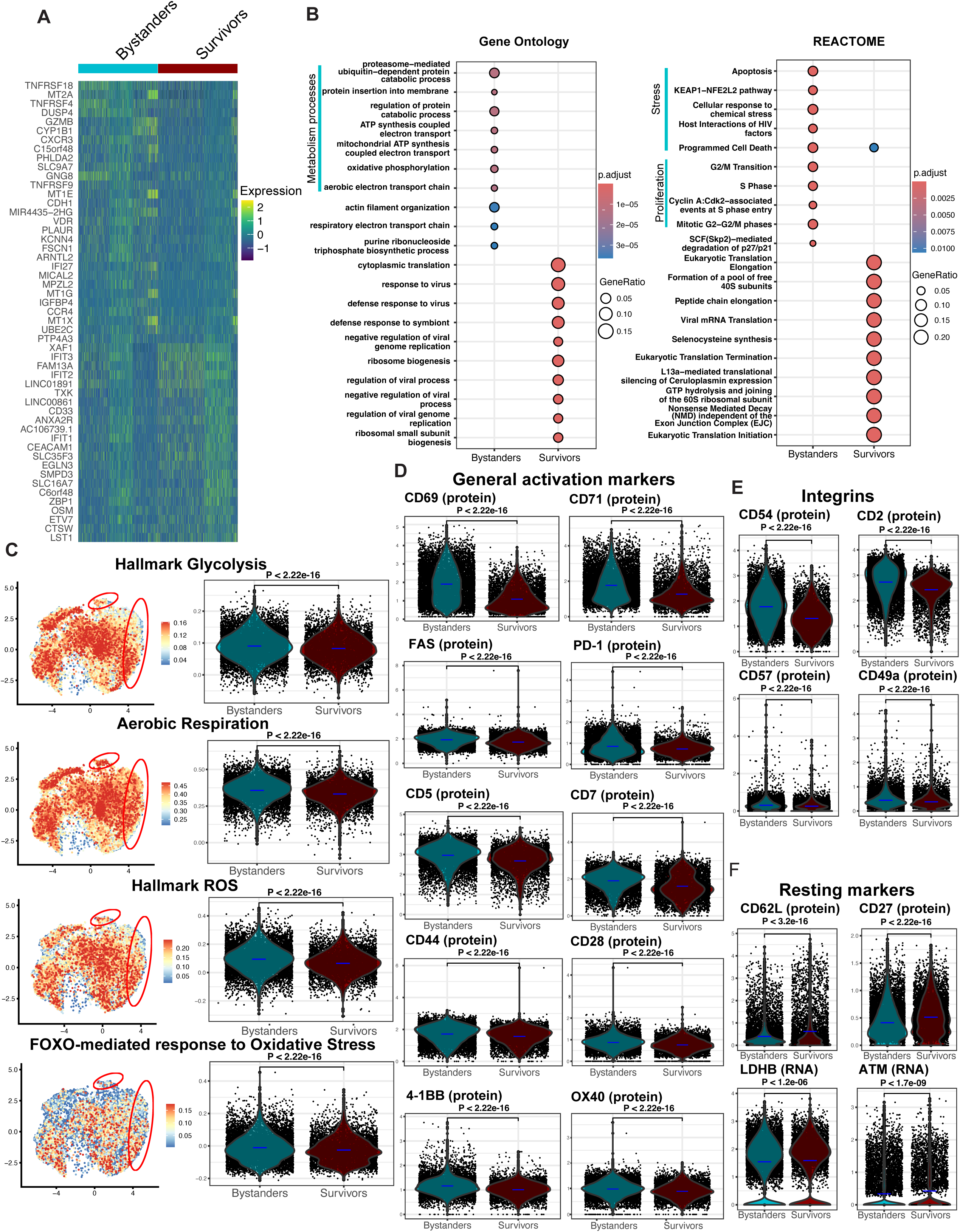
Quiescent metabolic and activation profiles are enriched in CTL-resistant infected cells. **(A)** Heatmap of top 30 differentially expressed genes across total pooled bystander and CTL-resistant survivor infected cells with a log_2_ fold-change difference of at least 0.1 passing min.pct ≥ 0.1 and P_adj_ < 0.05. **(B)** Top 10 differentially enriched Gene Ontology BP (left) and REACTOME (right) pathways from genes-sets passing filters defined in (**A**). **(C)** Module scoring of metabolism and oxidative stress gene-sets, mapped to WNN clusters. **(D)** ECCITE-seq-derived surface expression of selected surface protein activation markers, **(E)** integrins (activation-induced), and **(F)** resting. Expression of selected surface protein and gene-expression resting markers. For **(C)-(F)**, P-values of Wilcoxon tests are shown.

To assess if the immunophenotypes of CTL-resistant survivor cells followed the course of their transcriptional states, we quantified surface protein expression of markers of T-cell activation/differentiation and function in our ECCITE-seq data. Our analysis revealed 2-fold and 1.5-fold downregulation of the early activation markers CD69 and CD71 (the transferrin receptor) (Figure 3D). Similarly, other activation induced markers including Fas, PD-1, CD5, CD7, CD44 and the costimulatory receptors CD28, 4-1BB and OX40 were markedly down in survivors compared to bystander cells (Figure 3D). Activation induced integrins CD54 (ICAM), CD2, CD57 and CD49a, important for the stabilization of immune synapses with CTLs, were also robustly down in survivor cells (Figure 3E). Conversely, resting markers were upregulated in survivor cells compared to bystanders, including the adhesion molecule CD62-L (L-selectin) highly expressed in naïve and resting central memory T-cells (Figure 3F). These results highlight an enrichment of transcriptional programs denoting metabolic quiescence and redox balance, as well as a resting immunophenotypic state, as the most widely shared features of HIV-expressing cells that survive CTL attack.

### CTL-resistant, HIV-expressing cells have reduced levels of metabolic flux and oxidative stress

Since transcriptional and surface protein profiling of HIV-expressing survivor cells implicated quiescence and reduced oxidative stress, we next evaluated the bioenergetic demand and dependency of these cells via Single Cell ENergetIc metabolism by profilIng Translation inHibition (SCENITH)^44^. This method measures puromycin incorporation as a readout for new protein translation to measure the bioenergetic capacity of single cells through flow cytometry.

The use of different metabolic inhibitors targeting separate arms of ATP production allows for the deconvolution of glucose dependence (using OXPHOS inhibition) and mitochondrial dependence (using glycolysis inhibition). SCENITH analysis of HIV-expressing survivors revealed 1.22-fold and 1.25-fold lower levels of new protein translation compared to non-targeted bystander HIV-expressing cells in the same culture and HIV-expressing cells in the absence of CTL pressure, respectively (Figure 4A). This lower bioenergetic capacity in survivor cells was mainly attributed to reduced dependency on glucose oxidation, as survivors exhibited similar reductions in new protein translation when OXPHOS was inhibited, and did not show differences in translation compared to bystanders and infected cells without CTL pressure when glucose oxidation alone or both processes were inhibited (Figure 4B). These results confirmed that CTL-resistant, HIV-expressing survivors are enriched for cells with overall lower levels of metabolic flux compared to non-targeted HIV-expressing cells and that this reduced bioenergetic capacity is a result of lower glucose metabolism.

**Figure 4.**
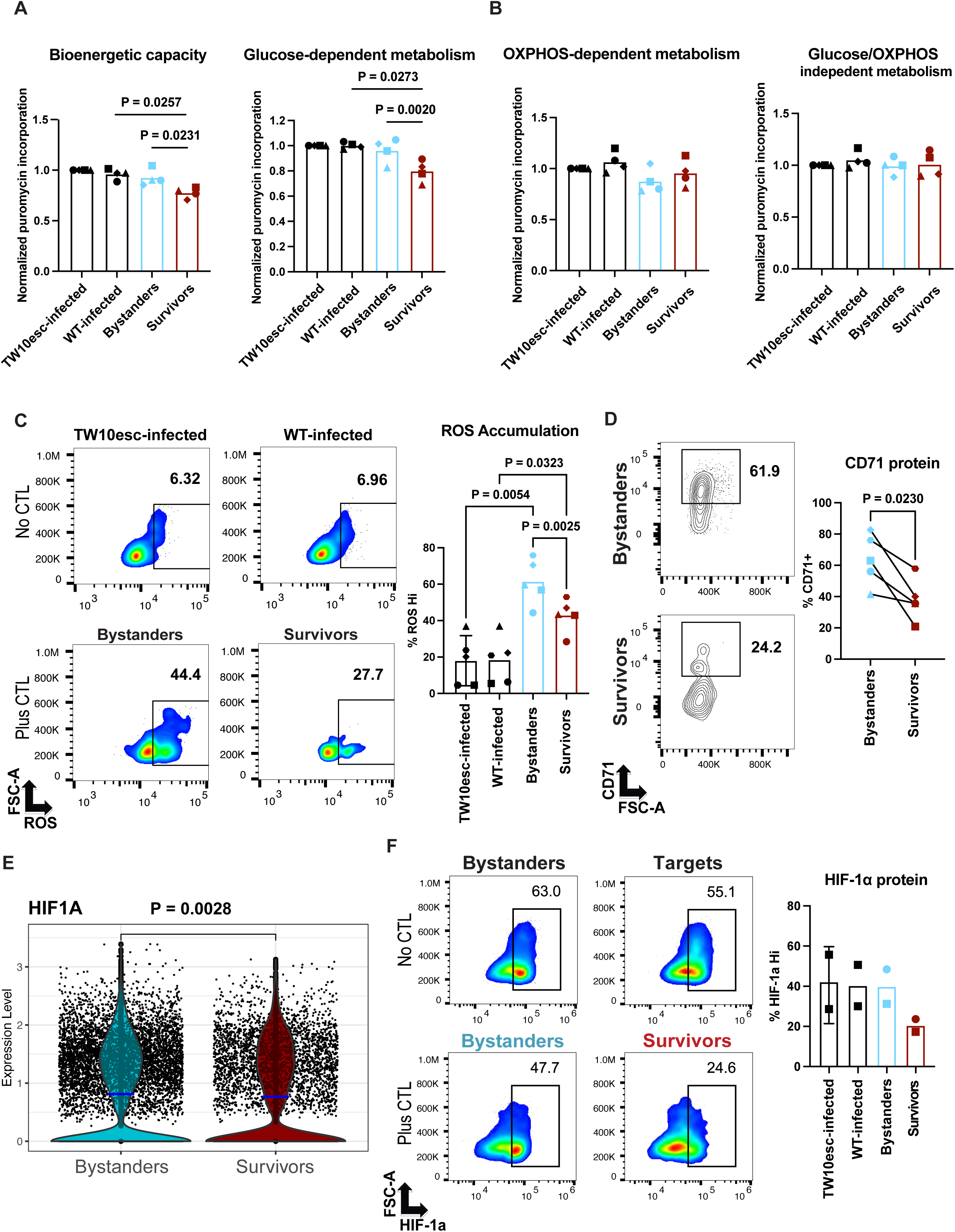
Reduced metabolic flux and oxidative stress in CTL-resistant infected cells. **(A)-(B)** SCENITH metabolic profiling of cells infected with either HIV_JRSCF TW10esc_ or WT HIV_JRSCF_ following either co-culture with an HIV-Gag-specific CTL clone or a parallel No CTL condition. In this assay, puromycin incorporation is used as a readout for metabolic activity in cells. Cells were separated into untreated and oligomycin A treated groups to measure total and glucose dependent metabolism respectively **(A)**, and 2-deoxyglucose treated and dual inhibitor treated groups to measure OXPHOS dependent metabolic activity and shut down both of these arms of metabolism respectively **(B). (C)** Quantification of intracellular ROS accumulation in the same populations as above using CellROX green or DCFH-DA. **(A)-(C)** Show results from 4 donors with P-values calculated by ANNOVA using Tukey’s multiple comparisons test. **(D)** Flow cytometry data showing surface CD71 expression in viable bystander and survivor cells surviving killing assays. **(E)** *HIF1A* transcript expression from previous ECCITE-seq of bystander and survivor cells. **(F)** Intracellular protein expression of HIF1-α in the same groups as **(A)-(C)**. For **(D)** P-value from a paired T-test is shown. For **(E)**, P-values of Wilcoxon test is shown.

Since our preliminary transcriptomics data indicated that CTL resistant survivors have dual downregulation of metabolic and oxidative stress gene-sets (Figures 3B and 3C), we reasoned that survivor cells may exhibit decreased intracellular ROS. To address this, we coupled our killing assay readouts with flow cytometry-based detection of intracellular ROS accumulation using CellROX green and DCFH_2_-DA^45,46^. These are compounds that become highly fluorescent when oxidized following reactions with a wide array of reactive oxygen species. Analysis of ROS accumulation in infected cells cultured with or without CTLs revealed that the presence of CTLs led to an increase in ROS accumulation in both survivor and bystander HIV-expressing cells compared to HIV-expressing cells cultured in the absence of CTLs (Figure 4C), suggesting that the culture microenvironment during a killing assay induces increases in intracellular ROS. Survivor cells, however, showed selection for cells with lower levels of ROS accumulation compared to bystanders in the same wells (Figure 4C), suggesting that ROS hi cells are preferentially susceptible to CTL-mediated death. Notably, survivors were also drastically depleted of cells expressing either CD69 or the transferrin receptor, CD71, responsible for intracellular uptake of iron (Figures 3D and 4D), implying that HIV-expressing cells with lesser activation states have a resistance advantage to CTL attack. An additional arm of cellular stress that culminates in the accumulation of intracellular ROS and metabolic reprogramming is the hypoxic response, which follows cellular activation in immune cells even in the presence of oxygen^47^. The Hallmark hypoxia gene-set as well as direct *HIF-1A* transcript levels from our sequencing analysis were found to be significantly reduced in CTL-resistant survivors compared to bystanders and these reduced levels were also confirmed for HIF1-A at the protein level (Figures 4E, 4F and S3D). Overall, these results demonstrated an enrichment towards metabolic quiescence and reduced oxidative stress in CTL-resistant, HIV-expressing cells. As CTL-derived granzyme A and granzyme B can both induce rapid mitochondrial depolarization and associated increases in intracellular ROS in targeted cells^48,49^, our results suggest that superior redox balance by a sub-population of HIV-expressing cells protects them from CTL-mediated elimination.

### Pharmacological induction of ROS and hypoxia sensitizes infected cells to CTL-mediated elimination

We next sought to address whether manipulating the redox state of HIV-expressing cells with the FDA-approved compound Deferoxamine (DFO) could improve CTL-mediated elimination of these cells. DFO is an iron chelator used for the treatment of excessive iron overload^50^ which stabilizes HIF-1a accumulation to induce a hypoxic mimic response and oxidative stress^51–54^. We observed that - in the absence of CTLs - treatment with DFO induced robust intracellular accumulation of ROS in infected cells (Figures 5A). This was complemented by a marked upregulation of CD71 surface expression among DFO-treated cells, an additional feature of CTL- sensitive infected cells (Figures 5B, S4A-B). Of note, DFO treatment also resulted in increased the frequencies of cells expressing HIV-Gag (Figure S4C). However, Despite the increase in initial HIV-expressing cells by DFO treatment before co-culture with CTLs, the same numbers of CTLs added across conditions were still able to eliminate a substantially greater proportion of HIV-expressing cells in the DFO-treated condition as compared to the vehicle control (DMSO), indicating a prominent role of DFO in sensitizing HIV-expressing cells to CTL elimination through mechanisms beyond latency reversal (Figures 5C and 5D). Thus, by reversing key metabolic features linked to CTL resistance, DFO potently sensitized HIV-expressing cells to improved CTL-mediated elimination. These results support the induction of ROS buildup in infected targets as a mechanism used by CTL to eliminate these cells and also highlight the ability of DFO to not only induce ROS but also additional arms of oxidative stress, such as the intracellular iron uptake transporter CD71, which can synergize to sensitize targeted cells to CTL-mediated elimination by engaging reactive oxygen- and reactive iron-based cell death pathways (Figure 5E)^48,49,55^.

**Figure 5.**
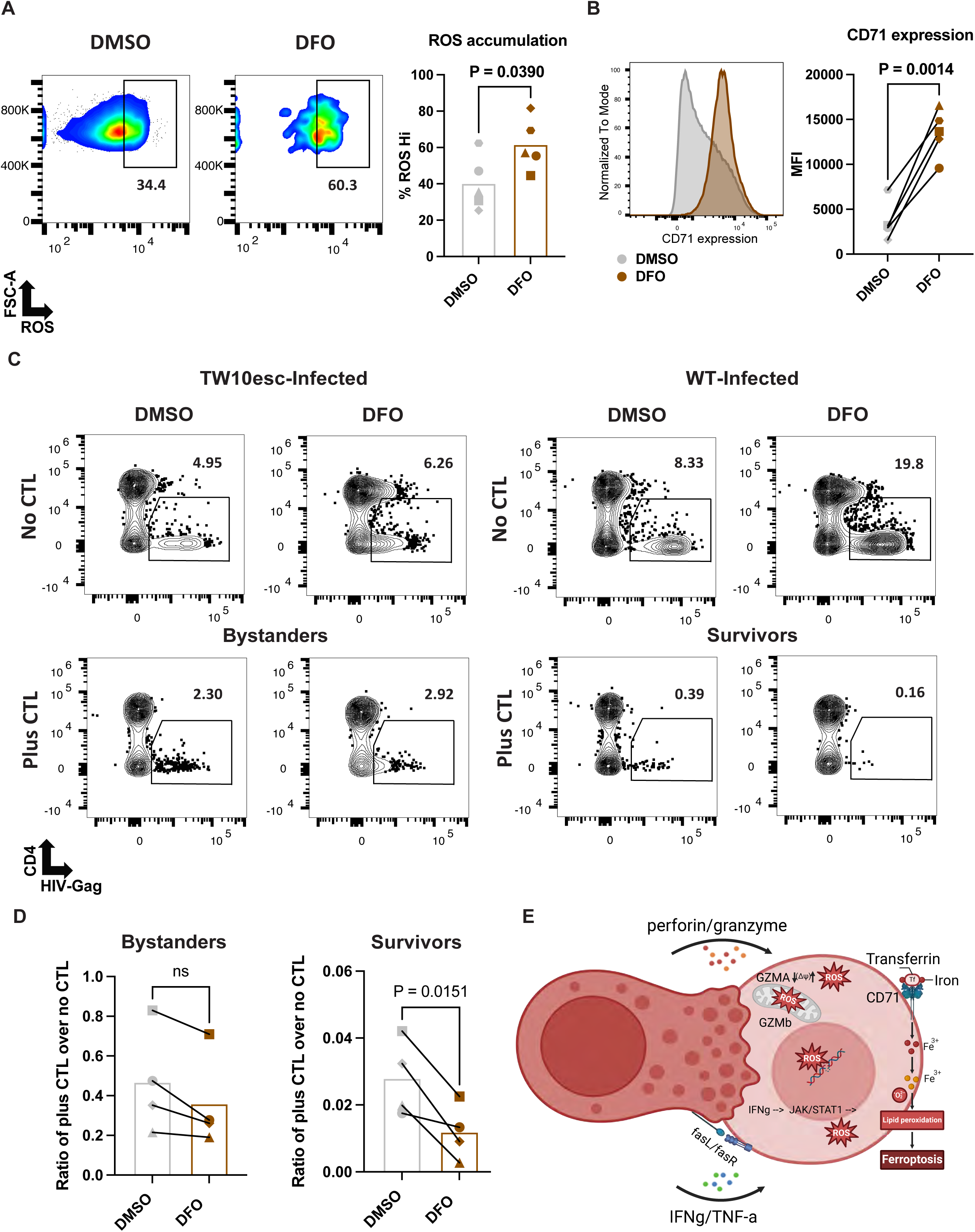
Induction of oxidative stress with Deferoxamine (DFO) sensitizes HIV-infected cells to CTL-mediated elimination. **(A)** Representative and summary (n=5 donors) flow cytometry plots showing intracellular ROS accumulation on HIV_JRSCF TW10esc_ or WT HIV_JRSCF_ infected cells following 72 hour treatments with either DFO or vehicle (DMSO). **(B)** Representative and summary (n=5 donors) flow cytometry plots measuring surface CD71 expression on HIV_JRSCF TW10esc_ or WT HIV_JRSCF_ infected cells following 72 hour treatments with either DFO or vehicle. **(C and D)** Representative (**C**) and summary (**D**) flow cytometry results depicting impact of DFO treatment on CTL-mediated elimination of infected cells. As in other experiments, cells infected with either WT HIV_JRCSF_ (targets/survivors) or escape mutant HIV_JRSCF_TW10esc_ (bystanders) were labeled with different cell-trace dyes and mixed together. Pooled cells were treated with DFO or DMSO (control) and then cultured with or without Gag-TW10-specific CTL. Shown are percentages of infected cells (CD4^-^, HIV-Gag**^+^**) following gating on cell-trace dyes. **(D)** Quantification of infected-cell killing as ratios of total viable cell counts from plus CTL conditions over no CTL conditions. **(E)** Schematic of proposed mechanisms of ROS-induced cell death mediated by CTL attack and improved by DFO treatment. For **(A), (B)** and **(D)** P-values from paired T-tests are shown.

### Reservoir-harboring cells that express HIV RNA exhibit CTL resistance metabolic features

Having identified and functionally validated a link between a transcriptional signature denoting quiescence and redox balance to CTL-resistance, we probed its *in vivo* relevance by asking whether this signature was present in cells harboring the HIV reservoir in ART-treated individuals. We leveraged a published data-set to focus on expanded T-cell clones containing cells found to be actively transcribing HIV RNA *ex vivo* reasoning that CTL resistance is most relevant to this ’active’ reservoir than to latently infected cells^22^. We scored expression of metabolic and oxidative stress modules in these cells and observed a marked downregulation of aerobic respiration, glycolysis, OXPHOS, hypoxia and ROS response gene-sets among HIV RNA**^+^** T-cell clones compared to HIV RNA- counterparts, highly reminiscent of the profiles of HIV-expressing CD4**^+^** T-cells that survived CTL attack *in vitro* (Figures 6A-6C). This suggests that the novel mechanism identified in the current study - of CTL resistance through metabolic quiescence- may contribute to the persistence of HIV-expressing CD4**^+^** T-cells *in vivo*.

**Figure 6.**
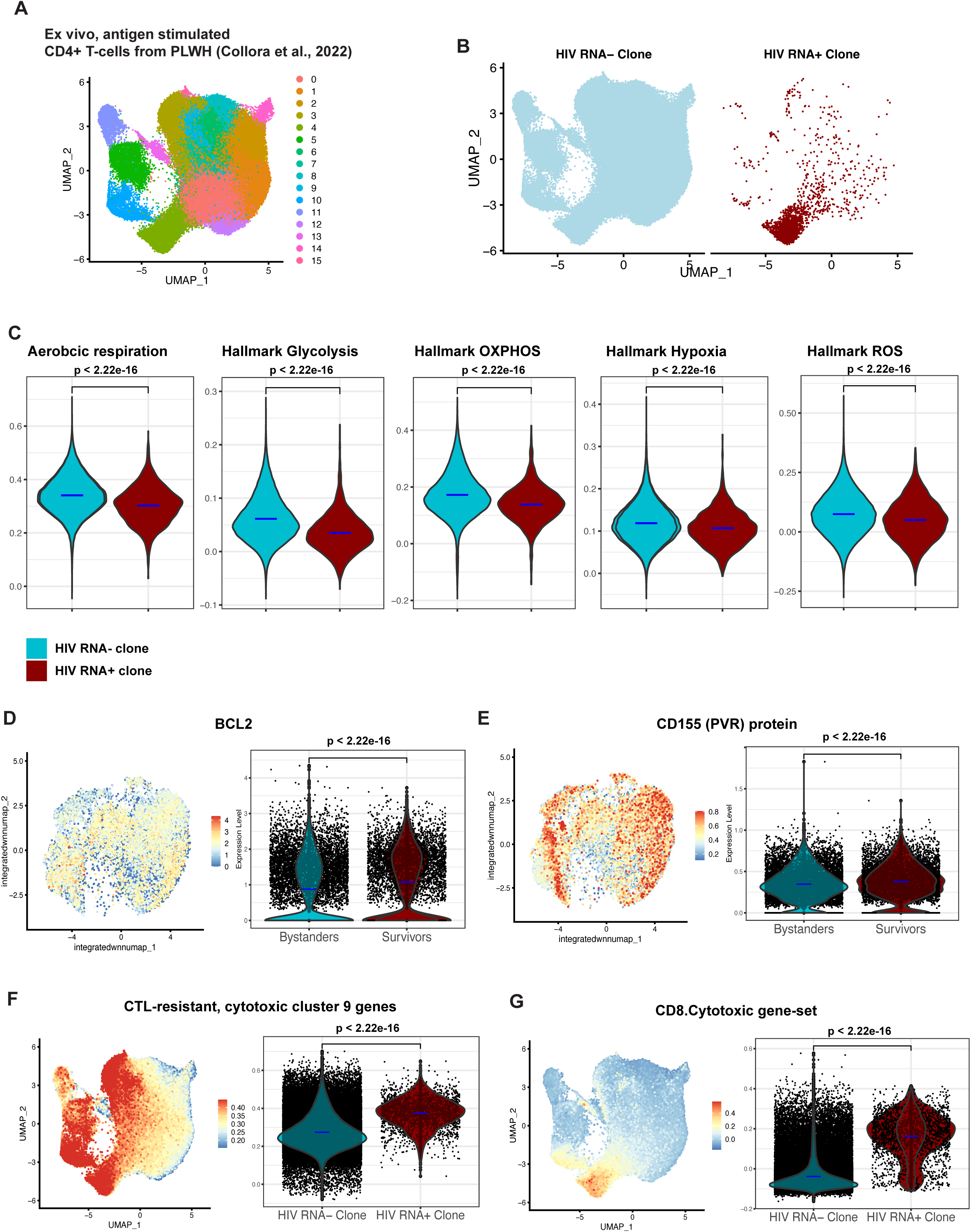
Molecular features of CTL-resistant infected cells are recapitulated in *ex vivo* reservoirs from PLWH. **(A)-(B)** Re-analysis of memory CD4**^+^** T-cells from PLWH in the antigen-stimulated data set from Collora *et al*., 2022^22^ showing unsupervised gene expression clustering of cells and labeling based on HIV RNA expression among T-cell clones. **(C)** Module scoring of metabolism and oxidative stress gene sets enriched in CTL-resistant survivors on the same cells from **(A)-(B)**. **(D)-(E)** UMAP feature plot and violin plots showing gene expression of *BCL2* and surface protein expression of CD155 PVR in bystander and survivor cells. **(F)** Module scoring of CTL-resistant WNN cluster 9 overexpressed genes on HIV RNA- and HIV RNA**^+^** T.-cell clones from **(A)-(B)**. **(E)** Module scoring of “CD8 cytotoxic” gene set from Szabo *et al*., 2019^36^ on HIV RNA- and HIV RNA**^+^**T-cell clones. All P-values shown are from a Wilcoxon test.

### Support for multiple additional mechanisms of resistance in HIV reservoirs

Advances in understanding the role and mechanisms of CTL-resistance in HIV persistence will be supported through bidirectional synergies between the profiles of CTL-resistant cells provided in the current study and ongoing advances in characterizing *in vivo* HIV reservoirs. Above, we have leveraged this framework to generate and functionally validate a hypothesis, which was then cross-referenced with profiles of *bona fide* reservoirs. Here, we reverse this directionality, examining whether previously proposed mechanisms of CTL resistance, identified through *ex vivo* observations, align with the functionally validated resistant cell profiles revealed by our dataset.

BCL-2 overexpression has been established as one mechanism by which HIV-infected CD4**^+^** T-cells can resist CTL and to date is the only validated therapeutic target to improve sensitization of infected cells to CTL elimination^26–28^. In the current study, *BCL2* transcripts were significantly overexpressed in CTL-resistant survivor cells compared to bystander cells with the highest mean expression level found in CTL-resistant cluster 4 (Figures 6D and S5A), further underscoring the relevance of this pathway in immune evasion.

Another proposed mechanism of resistance involves the immune checkpoint ligand PVR (CD155), which has been reported to be overexpressed on *ex vivo* CD4**^+^** T-cells harboring intact HIV proviruses. This overexpression was suggested to deliver a ‘don’t kill me signal’ to CTLs, promoting immune evasion. Consistent with this hypothesis, our findings show that PVR expression is elevated in CTL-resistant survivors relative to bystanders, with the highest levels found in cluster 11 (Figures 6E and S5B)^12^. This observation is consistent with the role of PVR as a factor contributing to CTL resistance in reservoir cells.

Multiple studies have reported that the HIV reservoirs that persist over time in ART-treated PLWH are preferentially harbored in CD4**^+^** T-cell clones with a cytotoxic profile^22,56^. This has led to the hypothesis that these cells may be equipped with mechanisms to resist cytotoxic effector molecules (e.g. perforin and granzyme) as a means of avoiding self-destruction^16^. We leveraged our survivor/bystander comparison to assess this by defining a gene module based on the overexpressed genes in CTL-resistant cytotoxic cluster 9 (Table S3) and applying this to *ex vivo* HIV RNA**^+^** and HIV RNA- CD4**^+^** T-cell clones from this previous study. In parallel we also reanalyzed this published data using the “CD8 Cytotoxicity” module from Szabo *et al*., 2019 that we had previously observed to be enriched in CTL-resistant cluster 9^22,36^. These analyses revealed a robust enrichment of both cytotoxic modules on HIV RNA**^+^**T-cell clones compared to HIV RNA- T-cell clones (Figures 6F-6G). This strong alignment between reservoir-harboring cells and the CTL-resistant cells in cluster 9 lends support to the hypothesis that resistance to killing contributes to the enrichment of HIV reservoirs in cytotoxic CD4**^+^**T-cells.

### Single-cell multi-omic profiling reveals complex relationship between MHC-I expression and CTL susceptibility

The HIV-Nef protein partially downregulates MHC-I downregulation on the surface of HIV-infected cells as a mechanism of partial immune evasion^57^. Thus, one would predict that cells with the lowest levels of surface MHC-I would be enriched our survivors. While this was observed when all cells were analyzed collectively (Figure S5C), further examination within the context of our WNN-defined clusters revealed a more nuanced landscape. Whereas CTL-resistant clusters 2, 4, and 11 have amongst the lowest levels of surface MHC-I, CTL resistant cluster 9 has amongst the highest. Conversely, the highly CTL-susceptible cluster 5 expresses amongst the lowest levels of MHC-I (Fig. S5D). Further, when all cells were divided into low (lower tercile), intermediate (middle tercile), and high (upper tercile) expressors of surface MHC-I, we observed that cells expressing intermediate and high levels of MHC-I were appreciably represented amongst CTL-resistant survivors (Fig. S5E and S5F). Similarly, while *BCL2* was overexpressed and PVR were under-expressed in collective survivors (Figure 6C-6D), cluster-specific analyses uncover extensive diversity in expression levels across clusters that do not necessarily track with overall CTL susceptibility of a given cluster (Figure S5A and S5B), further underscoring the diversity in resistance mechanisms across clusters. These findings reinforce the value of a single-cell multi-omic ’atlas’ of CTL resistance by further highlighting its ability to capture known and proposed mechanisms of CTL resistance / immune evasion, while revealing underlying complexities which will foster extensive follow-up studies.

## DISCUSSION

In this study, we provide direct evidence for the existence of populations of HIV-expressing primary CD4**^+^** T-cells that are capable of surviving rigorous attacks by HIV-specific CTLs. This phenomenon has been postulated previously in order to bridge gaps in the canonical model of HIV persistence - which attributes long-term survival of infected cells to viral latency - with recent observations indicating a degree of ongoing HIV expression and CD8**^+^**T-cell surveillance during ART^12,20–22,58^. However, to our knowledge, direct evidence that HIV-expressing cells can possess resistance to CTL has not previously been presented.

One important observation from our study is that cytotoxic CD4**^+^**T-cells that express HIV protein are relatively resistant to killing by CTL, compared to non-cytotoxic CD4**^+^** T-cells. The profiles of these cells show excellent agreement with cytotoxic CD4**^+^** T-cells that multiple studies have shown to disproportionately harbor HIV reservoirs in ART-treated PLWH, in particular those that continue to transcribe HIV^22,56^. Uncovering and counteracting the mechanisms by which these survive stringent CTL attack will require further study. Our results do suggest a potential role for somewhat diminished magnitudes of HIV transcription relative to non-cytotoxic cells, though this does not preclude roles for factors shown to protect other types of cytotoxic cells (e.g. CD8**^+^** T- cells and NK cells) from perforin/granzyme B mediated self-destruction, such as the granzyme B inhibitor SERPINB9^59^.

In addition to this cytotoxic profile, comprehensive multi-omic profiling and WNN clustering revealed that CTL resistant cells were enriched for resting phenotypes, and IFN response signatures, while CTL-sensitive cells exhibit high activation and proliferation. When all cells were considered collectively, metabolic quiescence and reduced intracellular ROS levels emerged as the most widely shared characteristic of CTL-resistant cells, and were validated as a novel mechanism of immune evasion by HIV-expressing cells. This was evident at the level of pathway enrichment analysis, but also at the level of numerous individual genes and proteins - such as dramatic selection amongst survivors against cells expressing CD69 and CD71. In tandem, multiple previously known or proposed mechanisms of CTL resistance were confirmed or supported by our dataset. These findings suggest that CTL-resistant cells evade immune clearance not through a single pathway but via distinct molecular and cellular programs. This is further supported by our observation that two established mechanisms of CTL resistance - MHC-I downregulation and *BCL2* overexpression - show the expected patterns of selection when all cells were analyzed collectively, but no consistent patterns at the levels of WNN-defined clusters of cells. We therefore propose that the datasets presented in this study serve as a foundational resource for advancing a comprehensive and nuanced understanding of CTL resistance in HIV.

To build upon this foundation towards therapeutic strategies, we linked transcriptional signatures of metabolic quiescence to low levels of the metabolic byproduct ROS, and demonstrated that the FDA-approved compound DFO markedly improves CTL-mediated elimination of HIV-infected cells. DFO was selected due to its ability to target multiple facets of resistance observed in survivor cells, by increasing CD71 surface expression and inducing hypoxia which culminates in increased intracellular ROS accumulation^51–54^. ROS exerts toxicity by interacting and damaging different essential macromolecules inside of the cell, including DNA^60^. Indeed, ROS buildup as a result of granzyme-mediated mitochondrial damage has been reported to contribute to cell death following a CTL hit^48,49^. However, ROS can also synergize with increased iron uptake via CD71 to induce the Fenton reaction and ROS/iron- mediated ferroptosis, a mechanism not yet studied in CTL-mediated clearance of infected cells^55^. Whether other compounds such as resveratrol or the FDA-approved drug atovaquone, which directly induce ROS production via disruption of the electron transport chain, can have similar or combinatorial effect with DFO will be of future interest^53,61–63^.

Our study points to further putative novel modalities of resistance for future exploration. Prominent amongst these was the observation of high levels of interferon stimulated genes amongst survivors - to some extent these overlapped with metabolically quiescent clusters, exemplifying that multiple features of resistance can overlap within populations of cells with similar states. One possibility is that these are cells sensing high levels of CTL-derived IFN-γ after surviving a failed killing attempt. Indeed, increased IFN-γ production by CTLs has been demonstrated in the context of poor and prolonged synapse formation with T-cell and macrophage targets and can have consequences on the inflammatory profile of the targets^64^. Alternatively, high levels of basal IFN responses in populations of HIV-infected CD4**^+^** T-cells may be a driver of resistance. For example, IFN responses can have direct effects on the metabolic states of cells that may impact susceptibility to CTL, such as through IFN-γ-mediated suppression of mTORC1 signaling and IFIT1- and IFIT3-mediated reductions in translation capacity^65,66^.

Importantly, the enrichment for metabolic quiescence and redox balance in CTL-resistant survivors also showed remarkable agreement with the metabolic signatures of rare reservoir-harboring cells studied *ex vivo* from PLWH, specifically with the subset transcribing HIV after antigenic stimulation. This convergence supports the clinical relevance of the mechanisms of CTL resistance defined by our study and provides rationale for novel therapeutic approaches to address HIV persistence. Our observations may also have translational implications outside of the context of HIV. In the context of T-cell lymphoma, enrichment of quiescent signatures has also been observed in metastatic cells that disseminate away from the primary site of malignant transformation, suggesting that immune evasion through immune quiescence may contribute to other pathologies where persistent pathologic CD4**^+^** T-cells are targeted^34,67^. Collectively, we provide an extensive atlas for cross-referencing cellular programs that may affect target cell elimination by CTL and propose metabolic perturbation of reservoir harboring cells as a novel therapeutic approach to advance strategies towards an HIV cure.

### Limitations of the Study

A limitation of our study is that the full diversity of CD4**^+^**T-cell differentiation and polarization states is unlikely to be reflected in our cultured T_CM_ target cells. While aiming to generate relatively homogenous target cells was a strategic decision made to increase our ability to uncover core functional drivers of CTL resistance, it may be important to extend this to other target populations - for example, CD4**^+^** T-cells that are intentionally polarized towards T follicular helper or cytotoxic profiles. Some degree of this complexity did, however, emerge in our system - notably, cytotoxic CD4**^+^** T-cells with transcriptional profiles that closely aligned with *ex vivo* reservoir-harboring cytotoxic CD4**^+^**T-cells. There are also limitations in the degree to which encounters with CTL and resulting selection *in vitro* stand-in for those that occur *in vivo* and across various tissues. We stress the importance of identifying convergences between features of survivor cells selected in our system and *bona fide ex vivo* reservoir harboring cells in order to discriminate targets with therapeutic potential.

## Acknowledgments

We thank all study participants who devoted time to our research. The following reagents were obtained from the NIH HIV Reagent Program, Division of AIDS, NIAID, NIH: Anti-Human Immunodeficiency Virus (HIV)-1 gp120 Monoclonal Antibody (PGT121) cat # ARP-12343, anti-Human Immunodeficiency Virus (HIV)-1 gp160 Monoclonal Antibody (N6/PGDM1400x10E8) cat # ARP-13390, and JR-CSF Infectious Molecular Clone (pYK-JRCSF) cat # ARP-2708). We acknowledge the Biological Resources Branch Preclinical Repository, National Cancer Institute, for providing IL-2 and IL-15. Research reported in this publication was supported by the National Institutes of Allergy and Infectious Diseases of the National Institutes of Health under Award Numbers UM1AI164565 (R.B.J), R01AI181626 (R.B.J), R01AI176601 (R.B.J), R01AI170245 (R.B.J), R01AI165031 (R.B.J), UM1AI164562 (R.B.J), R21AI170246 (R.B.J), and R01AI147845 (R.B.J). UM1AI164565 and UM1AI164562 were also supported by the National Institutes of Diabetes and Digestive and Kidney Diseases, the National Institute of Mental Health, the National Institute of Neurological Disorders and Stroke, the National Institute on Drug Abuse, and the National Heart, Lung, and Blood Institute. The content is solely the responsibility of the authors and does not necessarily represent the official views of the National Institutes of Health.

## Author contributions

A.H, L.L and R.B.J. initiated the project and were responsible for experiment design. A.H and L.L performed the majority of experiments with assistance from J.W., T.T.H., and F.W. in specific assays; M.C. and C.K. were directly involved in patient recruitment, sample collection and study design; A.H., P.Z. and M.A.A. performed the bioinformatic analyses with input from D.B. and R.B.J.; A.H., L.L., and R.B.J. wrote the paper with all authors providing input on the manuscript.

## Declaration of generative AI and AI-assisted technologies

During the preparation of this work, the author(s) used GPT 4.0/OpenAI in order to obtain feedback on and refine writing structure and style. After using this tool or service, the author(s) reviewed and edited the content as needed and take(s) full responsibility for the content of the publication.

## Supplemental information

Table S1. Related to figure 1G-1F.

Table S2. Related to figure 3A.

Table S3. Related to figure 6F.

## STAR Methods

### KEY RESOURCES TABLE

| REAGENT or RESOURCE | SOURCE | IDENTIFIER |
| --- | --- | --- |
| **Antibodies** |  |  |
| CD4 BV785 clone OKT4 | Biolegend | Cat. #317442 |
| CD4 AF700 clone OKT4 | Biolegend | Cat. #317426 |
| CD3 BV785 clone OKT3 | Biolegend | Cat. #317330 |
| CD8 BV605 clone SK1 | Biolegend | Cat. #344742 |
| CD107a APC/Cyanine 7 clone H4A3 | Biolegend | Cat. #328630 |
| CD107a PE clone H4A3 | Biolegend | Cat. #328608 |
| CD69 BV650 clone FN50 | Biolegend | Cat. # 310934 |
| CD71 Pe/Cyanine 7 clone CY1G4 | Biolegend | Cat. # 334112 |
| HLA-A,B,C PE clone W6/32 | Biolegend | Cat. # 311406 |
| IL-17 BV605 clone TC11-18H10.1 | Biolegend | Cat. # 512325 |
| IL-4 PE/Dazzle™ 594 clone MP4-25D2 | Biolegend | Cat. # 500831 |
| IFN-γ Pe/Cyanine 7 clone B27 | Biolegend | Cat. # 506518 |
| IL-22 PerCP/Cyanine5.5 clone 2G12A41 | Biolegend | Cat. # 366709 |
| TotalSeq-C Human Universal Cocktail V1.0 | Biolegend | Cat. # 399905 |
| TotalSeq™-C0251 anti-human Hashtag 1 clones LNH-94; 2M2 | Biolegend | Cat. # 394661 |
| TotalSeq™-C0251 anti-human Hashtag 3 clones LNH-94; 2M2 | Biolegend | Cat. # 394605 |
| TotalSeq™-C0251 anti-human Hashtag 4 clones LNH-94; 2M2 | Biolegend | Cat. # 394607 |
| HIF1- $\alpha$ AF488 clone H1alpha67 | Novus Biologicals | Cat. # NB100-105F |
| Zenon Human IgG labeling Kit AF647 | ThermoFisher | Cat. #Z25408 |
| Zenon Human IgG labeling Kit R-PE | ThermoFisher | Cat. #Z25455 |
| N6/PGDM1400x10E8 | BEI resources | ARP-13390 |
| PGT121 | BEI resources | ARP-12343 |
| Ultra-LEAF™ Purified anti-human CD3 Antibody | Biolegend | Cat. #317325 |
| Ultra-LEAF™ Purified anti-human CD28 Antibody | Biolegend | Cat. #302933 |
| **Bacterial and virus strains** |  |  |
| JRCSF | BEI resources | ARP-394 |
| JRCSF TW10escape T242N mutant | Genescript | U1998GG250 |
| JRCSF SL9escape mutant | Genescript | U423KGJ130 |
| **Biological samples** |  |  |
| Human peripheral blood leukopak fresh HLA A02 B57 | Stemcell | Cat. #70500 |
| Human blood buffy coat | the Gulf Coast<br>Regional Blood Center |  |
| **Chemicals, peptides, and recombinant proteins** |  |  |
| TECIN (Teceleukin) Recombinant Human IL2 (rIL-2) | NCI | C166111-01 |
| Recombinant Human IL-15 | NCI | NSC. #745101 |
| Dynabeads™ Human T-Activator CD3/CD28 for T Cell Expansion and Activation | ThermoFisher | Cat. #11131D |
| Anti-Human IL-12 | Peptotech | Peptotech 500-P154G |
| Anti-Human IL-4 | Peptotech | Peptotech 500-P24 |
| Recombinant Human TGF-β1 | Peptotech | Peptotech 100-21 |
| Cell trace far red (CTFR) | ThermoFisher | Cat. #C34564 |
| Cell trace CFSE | ThermoFisher | Cat. #C34570 |
| Cell trace Violet (CTV) | ThermoFisher | Cat. #C34557 |
| Cell trace Blue (CTB) | ThermoFisher | Cat. #C34574 |
| Live/Dead Fixable Aqua dead cell stain kit | ThermoFisher | Cat. #L34957 |
| **Critical commercial assays** |  |  |
| EasySep™ Human Naïve CD4<sup>+</sup> T Cell Isolation Kit | Stemcell | Cat. #19555 |
| EasySep Human CD4 Positive Selection Kit II | Stemcell | Cat. #17852 |
| EasySep Human CD8 Positive Selection Kit II | Stemcell | Cat. #17853 |
| **Deposited data** |  |  |
| GEO accession number pending |  |  |
| **Experimental models: Cell lines** |  |  |
| HEK 293T Cells | ATCC |  |
| **Software and algorithms** |  |  |
| R version 4.3.2 | R Core Team (2024) |  |
| FlowJo version 10.10.0 | BD Biosciences |  |

## METHOD DETAILS

### Participants and study approval

Study participants living with HIV were recruited from Whitman Walker Health (Washington, D.C). Leukapheresis samples from deidentified HIV negative individuals were obtained from Stemcell Technologies. Ethical approval to conduct this study was obtained from George Washington University and the Weill Cornell Medicine Institutional Review Boards (Weill Cornel Medicine #1805019252 and George Washington University FWA#00005945). All participants were adults and provided written informed consent.

### HIV infection model in cultured central memory CD4^+^ T cells (T_CM_)

The infection model was adapted from a previously described HIV latency model^30,31^. Peripheral blood mononuclear cells (PBMCs) were isolated from leukapheresis samples from HIV negative donors obtained from Stemcell technologies. Naïve CD4^+^ T cells were magnetically isolated from thawed PBMCs using the EasySep™ Human Naïve CD4**^+^** T Cell Isolation Kit (Stemcell) according to the manufacturer’s instructions. Purified naïve CD4^+^ T cells were then activated using human anti-CD3/CD28-coated magnetic beads (one bead for two cells, Gibco Cat. #11131D) in the presence of human anti-IL-4 (1µg/10^6^ cells Peprotech 500-P24), human anti-IL-12 (2µg/10^6^ cells Peprotech 500-P154G), and Tumor Growth Factor (TGF)-β1 (10ng/10^6^ Peprotech 100-21) to prevent cell polarization. After 3 days, cells were maintained at a concentration of 10^6^ cells /mL in media containing 50 IU of human recombinant IL-2. At day 7, the cultured T_CM_ were infected with either WT HIV_JRCSF_ or HIV_JRCSF TW10escape_. One quarter of the cells were infected by the addition of 200 units of virus (see virus production details below for unit description) per 1x10^7^ cells in 1mL and centrifugation at 1500xg for 2h at 37°C. Following this ’spinoculation’, the infected cell cultures were mixed in flasks containing the remaining three quarters of the cells at 2x10^6^ cells/mL in media containing 50 IU of human recombinant IL-2. At day 10, the cells were crowded in 96-well round bottom plates at a density of 2x10^5^ cells/200uL/well to enhance cell-to-cell transmission of HIV. On day 13, cells were cryopreserved in FBS with 10% DMSO and stored in liquid nitrogen for future experiments.

### Virus production and infections

JRCSF WT and JRCSF TW10esc plasmids were obtained from BEI-resources and Genscript, respectively. Plasmids were expanded using Stlb3 competent cells and maxi-prep from Qiagen (QIAGEN Plasmid Maxi Kit (25)). Plasmids were transfected into HEK 293T cells using Optimem media and FuGENE® 6 Transfection Reagent (Promega). Media was exchanged 24h after transfection for DMEM with 10% FBS. Two days following the media change, supernatants were passed through 0.45μm filters. Viruses were then concentrated overnight at 4°C using the PEG-it virus precipitation solution (System Biosciences) following the manufacturer’s instructions. Finally, the concentrated virus was frozen in aliquots of 200 µl. Across multiple titration experiments, 200 µl of virus for 1 x 10^7^ cells gave the most optimal results balancing infection efficiency and viability of productively infected cells.

### Isolation and expansion of HIV specific Cytotoxic T- lymphocytes (CTLs)

CTL specific for HIV epitopes Gag-TW10 and Gag-SL9 were isolated from PBMCs of 3 donors living with HIV. Briefly, 10 x 10^6^ PBMCs were peptide pulsed with HIV optimal peptides TW10 (TSTLQEQIGW, Gag residues 240-249) or SL9 (SLYNTVATL Gag residues 77-85) at 1μg/mL for 16h at 37°C in XVIVO-15 media. T cells that responded to the peptide by secreting IFN-γ were isolated using an IFN-γ secretion assay – Cell Enrichment and Detection Kit (PE) (Miltenyi Cat#130-054-202). The cells were then serially diluted and stimulated with PBMCs from HIV-negative donors that had been irradiated with 5,000 Rad, along with soluble anti-CD3/anti-CD28 in media containing 50IU IL-2 and 0.51ng/mL IL-15. Wells for selected from dilutions in which <1/5 wells showed growth, giving confidence that the cells in each well were derived from a single clone. CTL specificities were assessed by flow cytometry by assessing degranulation (CD107a surface exposure) upon the addition of 1μg/mL of the respective peptide.

### Killing assays and sequential killing assays

Infected cells from the previously described HIV Infection Model in cultured T_CM_ were thawed and rested for 2 days at 2 x 10^6^ cells per mL in media containing 50 IU of human IL-2. Prior to co-culture with CTL, cells expressing high levels of CD4 were depleted by magnetic selection using the EasySep™ Human CD4 Positive Selection Kit II (Stemcell); this step enriches for CD4^-^ infected cells (since HIV-infected cells downregulate CD4). Cells infected with WT HIV_JRCSF_ were labeled with cell trace far red (CTFR) (dilution 1:20,000) and cells infected with HIV_JRCSF TW10escape_ were labeled with cell trace carboxyfluorescein succinimidyl ester (CFSE) (dilution 1:5,000). CTFR-labeled WT HIV_JRCSF_-infected cells were mixed with CFSE-labeled HIV_JRCSF TW10escape_- infected cells at a ratio of 2:1 and 6 x 10^5^ cells from these mixed populations were aliquoted into wells of a 96-well plate. 3 x 10^5^ CTL clones specific for the HIV-Gag epitope TW10 ’TSTLQEQIGW’ were labeled with CellTrace™ Violet (CTV) (dilution 1:10,000) and were added to each well at a 1:2 effector:target ratio along with anti-CD107a APC-Cy7 (dilution 1:200). Killing assays were performed in R10 (RPMI 1640 supplemented with 10% FBS and 1% penicillin/streptomycin, 1% HEPES buffer, 5 mM L-Glutamine), with 50 IU of human recombinant IL-2 and 0.51 ng/mL of human recombinant IL-15 for 16h at 37°C. For Sequential Killing Assays, CTL were then removed by magnetic selection using the EasySep™ Human CD8 Positive Selection Kit II (Stemcell) and a fresh batch of CTL clone specific for the HIV-Gag epitope SL9 ’SLYNTVATL’ were added for the second round of killing, which was again for 16h at 37°C (no CTL control conditions were maintained in parallel). Post-second killing assay, the levels of infected cells were assessed by flow cytometry, staining extracellularly with anti-CD3 (OKT3, Biolegend), anti-CD4 (OKT4, Biolegend), anti-CD8 (SK1, Biolegend) antibodies and Live/Dead fixable Aqua stain (Thermofisher). Cells were then fixed with the BD Cytofix/Cytoperm™ Fixation/Permeabilization Kit according to the manufacturer’s instructions and intracellularly stained using an anti-HIV-Gag antibody (KC57, Beckman Coulter) before analysis on an Attune NxT flow cytometer.

### Sorting for ’survivors’ and ’bystanders’ following killing assays

Killing assays were set up as described above, with minor differences. For these sorting assays, cells infected with WT HIV_JRCSF_ were labeled with CFSE (dilution 1:5,000) and cells infected with HIV_JRCSF TW10escape_ were labeled with CTV (dilution 1:10,000). These were then mixed together before adding a TW10-specific CTL that had been labeled with CellTrace™ Blue CTB (dilution 1:10,000). No CTL control conditions were cultured in parallel. Post-killing, cells were harvested for staining with anti-CD4 BV785 (1:100), Live/Dead fixable Aqua (1:200), bNAbs PGT121 conjugated with Zenon Kit PE (1µg per 1 x 10^6^ cells) and bNAbs N6/PGDM1400x10E8 conjugated with Zenon Kit AF647 (1µg per 1 x 10^6^ cells) at 37°C for 1h. Live cells productively infected CFSE- labeled WT HIV_JRCSF_ and CTV-labeled HIV_JRCSF TW10escape_ were sorted by gating on the double positive population for PGT121 and N6/PGDM1400x10E8.

### Flow cytometry for SCENITH metabolic profiling

Infected bystander and target/survivor cells after a ‘killing assay’ with or without the addition of CTL were treated in R10 media as previously described^44^, with a few modifications. Briefly, cells were treated for 60 minutes with DMSO or with the following metabolic inhibitors - 2-Deoxy-D- Glucose (DG, final concentration 100mM), Oligomycin (Oligo, final concentration 1μM), or a combination of the drugs at the same final concentrations. Puromycin (final concentration 10μg/ml) was added during the last 30 minutes of the treatments with the metabolic inhibitors. Single-cell suspensions were stained for surface markers using the following antibodies anti-CD4 (OKT4, Biolegend), anti-CD8 (SK1, Biolegend), anti-CD7 (CD7-6B7, Biolegend), anti-CD71 (CY1G4, biolegend), anti-CD69 (FN50, Biolegend), anti-CD45RA (HI100, Biolegend) and anti-HLA-A-B-C (W6/32, Biolegend) along with a fixable live/dead stain (Thermofisher), in PBS for 30 minutes at 37°C. Cells were washed and fixed with 1X eBioscience™ FoxP3/transcription factor fixation/permeabilization solution (Thermofisher) for 45 minutes at 4°C. This fixation solution was washed away and cells were stained intracellularly with anti-HIV-Gag-p24 (KC57, Beckman Coulter), anti-puromycin (2A4, Biolegend) and HIF1-α (H1alpha67, Novus Biologicals) antibodies in 1X FoxP3/transcription factor permeabilization solution (Thermofisher) for 60 minutes at 37°C. Cells were washed one more time before analysis on an Attune NxT flow cytometry instrument.

### Flow cytometry for ROS accumulation

Infected bystander and target cells after a ‘killing assay’, with or without the addition of CTL, were treated with 500 nM CellROX green (Thermofisher) or 0.2X DCFH-DA (CELL BIOLABS) for 1 hour. CellROX solution was washed and single cell suspensions were stained using surface target antibodies against CD4 (OKT4, Biolegend), CD8 (SK1, Biolegend), CD7 (CD7-6B7, Biolegend), CD71 (CY1G4, biolegend), CD69 (FN50, Biolegend), CD45RA (HI100, Biolegend) and HLA-A-B-C (W6/32, Biolegend) along with a fixable live/dead stain (Thermofisher) in PBS for 30 minutes at 37°C. Cells were washed and then fixed with 1X Cytofix/Cytoperm™ solution (BD) for 20 minutes at room temperature. Fixation solutions were washed away and cells were stained intracellularly with anti-HIV-Gag-p24 (KC57, Beckman Coulter) antibody in 1X Perm/Wash™ (BD) for 30 minutes at 37°C. Cells were washed one more time before analysis on an Attune NxT flow cytometry instrument.

### ECCITE-seq compatible with a 10X genomics instrument

Cells were stained with DNA oligo-barcoded antibodies and loaded onto the 10x Chromium single-cell workflow (10X Genomics) as previously described^34^ with a few modifications. Approximately 0.05-0.3 x 10^6^ cells per sample were resuspended in R10 media and incubated with the TotalSeq™-C Human Universal Cocktail (Biolegend) for 30 minutes at 4°C. In some cases, TotalSeq™-C hashing antibodies #1, #3 and #4 (Biolegend) were spiked into individual samples, which allowed for multiplexing of survivor, bystanders and cells with no CTL into the same 10X lane. After staining, cells were washed 3 times in R10 media, followed by centrifugation (400 × g for 5 minutes at 4°C) and supernatant aspiration. After the final wash, cells were resuspended in PBS and filtered through 70-µm cell strainers. Stained cells from each sample were pooled and loaded into the 10x Chromium Single Cell Immune Profiling workflow (CG000330, revision G). Libraries were generated according to the 10x Chromium Single Cell Immune Profiling workflow instructions, pooled and sequenced on Illumina NovaSeq 6000 or a NovaSeq X.

### Single-cell data alignment

FASTQ files from the 10x gene expression, surface protein and TCR α/β libraries were processed using Cell Ranger v7.0.1 (10x Genomics), using either the count or the multi pipeline (when TCR libraries were available) with default parameters.

RNA transcripts from ECCITE-seq were mapped to a modified version of Cell Ranger’s GRCh38- 2020-A reference package (GRCh38 / GENCODE v32 annotations), with the addition of the HIV- 1 HXB2 reference sequence. TCR α/β libraries were mapped to Cell Ranger’s cellranger-vdj- GRCh38-alts-ensembl-7.1.0 reference package.

### Initial cell/gene level filtering

For cell filtering, cell barcodes with <200, >8,000 genes expressed or > 10% mitochondrial genes expressed were removed. For gene-level filtering, genes were filtered such that only those expressed at a level of <200 UMI or in < 3 cells were removed.

These cell- and gene-level filtered gene/barcode matrices were used for downstream analysis.

### Seurat normalization

The count matrices of all modalities were loaded into a Seurat v5 object (Seurat version 5.1.0). For gene expression (GEX) data, we normalized the count data using LogNormalize and scaled the resulting normalized values using ScaleData. For all other modalities (ADT, HTO, TCR α/β), we normalized the count data using centered-log ratio (margin = 2) and scaled the resulting normalized values using ScaleData.

### Hashing antibody demultiplexing

To separate individual samples from pooled cell libraries, samples were demultiplexed by their HTO using Seurat HTODemux with the positive quantile parameter set to 0.999.

### Harmony integration of GEX and ADT modalities across donors

To integrate datasets from different donors across separate 10X runs, datasets from each donor were first merged into one object using the merge Seurat function and then we used RunHarmony as previously described on GEC and ADT modalities to remove dataset-of-origin cell embeddings using the “batch” covariate, in this case “donor” included in the objects metadata^68^. Cells were then given probabilistic cluster membership based on K-means cluster centroids localization for each modality.

### Multimodal WNN analysis

We applied weighted-nearest neighbor (WNN) analysis on our ECCITE-seq data, enabling the integrative analysis of GEX and ADT modalities in the same cells as previously described^20^. Briefly, k-nearest neighbors graphs (k = 20) were first constructed, and the neighbors of each cell in each modality were identified using FindMultiModalNeighbors in Seurat V5. For graph construction, the first 15 principal components (PCs) of the log-normalized values from the GEX modality and the first 12 PCs of the centered-log ratio values from the ADT modality were used. The dimensionally reduced molecular profiles of the k-nearest neighbors of each cell were then averaged and compared with the actual values to obtain the residuals in an RNA-RNA, protein- protein and RNA-protein manner. Residuals were converted into affinity-based similarities using UMAP. Ratios of the within-modality and cross-modality affinities were softmax transformed to obtain the cell-specific modality weights. The weighted similarity then formed the basis of the construction of the WNN graph visualized on a WNN integrated UMAP. Clusters were called using the FindClusters function at a resolution of 0.5.

### Additional cell filters

WNN clusters 3 and 10 were filtered out after initial analysis due to disproportionately high expression of MT-genes compared to other clusters. Cells having > 0 normalized expression of *CD8A or CD8B* transcripts or > 0.5 normalized expression of CD8 protein were filtered out from further analysis.

### Differential gene expression and pathway enrichment analysis

Differential gene expression analysis across WNN clusters was performed using FindAllMarkers. The logfc.threshold (minimum log_2_ fold change) was set to 0.25 and min.pct (minimum proportion of cells expressing the given gene) was set to 0.25 to avoid inclusion of poorly expressed genes that can skew differential gene expression results. Only genes with an adjusted P-value of < 0.05 were further considered. Row normalized Z-scores from using scale.data were used for visualization on the heatmap focusing on the top 10 differentially expressed genes.

Genes passing the above filters were used for pathway enrichment analysis. ClusterProfiler was used to profile gene clusters on Gene Ontology terms. Specifically the biological process (GO:BP) subontology was used for enrichment analysis. Gene ontology terms were considered significant if the FDR-adjusted P values was < 0.05.

Similarly, total bystander and survivor populations were compared using FindAllMarkers with the logfc.threshold set to 0.1 and min.pct set to 0.1. Top 30 differentially expressed genes were used for visualization on the heatmap showing row-normalized Z-scores. Genes passing the logfc.threshold and min.pct filters were used for pathway enrichment analysis with ClusterProfiler as above but using both the GO:BP subontology as well as Reactome for gene cluster profiling and visualization.

### Module scoring on bystander and survivor cells from T_CM_-infection model

The AddModuleScore function of Seurat V5 was used for scoring gene expression modules. For Hallmark and Gene Ontology gene-sets, the msigdbr package was used to pull selected gene- sets from the Molecular Signatures Database (MSigDB).

T-cell activation and functionality gene-sets were pulled from Szabo *et* al., 2019^36^. The cytotoxic WNN cluster 9 gene-set was constructed from the results of differential gene expression analysis comparing WNN cluster 9 to all other clusters after all unwanted cell removal filter steps using the FindMarkers function and setting min.pct = 0.1 and logfc.threshold = 0.05. Genes were then filtered for those significantly overexpressed by WNN cluster 9 subletting on those with log_2_ fold change > 0.05 and adjusted P-value < 0.05.

### Module scoring on HIV RNA^+^ and HIV RNA- T-cell clones from PLWH

Single cell transcriptional and V(D)J reconstruction data from *ex vivo* antigen stimulated CD4**^+^** T- cell clones from PLWH was pulled from Collora *et al*., 2022^22^. HIV RNA**^+^** T cell clone cells were defined as 2 or more cells expressing the same CDR3 having at least one HIV-1 RNA**^+^** cell. HIV RNA- T-cell clones were defined as 2 or more cells expressing the same CDR3 with no member expressing HIV transcripts. As above, the AddModuleScore function of Seurat V5 was used for scoring gene expression modules of Hallmark and Gene On The AddModuleScore function of Seurat V5 was used for scoring gene expression modules of Hallmark and Gene Ontology gene- sets from MSigDB using msigdbr, the “CD8 cytotoxic” functional state from Szabo *et* al., 2019^36^, and the WNN cluster 9 overexpressed gene-set

## RESOURCE AVAILABILITY

### Data and code availability

ECCITE-seq data is being deposited to GEO. The accession number will be listed in the key resources table. This paper does not report original code.

## SUPPLEMENTARY FIGURE LEGENDS

**Supplementary Figure 1**: **(A)** Schematic of the HIV Infection Model in primary central memory CD4^+^ T cells. **(B)** Single nucleotide mutation in TW10 epitope of JRCSF converting threonine (T) to asparagine (N) to prevent binding of TW10 antigen to MHC-I. **(C)** Phenotype of HIV infection model by flow cytometry looking at CD45RA, MHC-I and HIV-Gag**^+^**from naïve CD4^+^ T cells at day 0 to infected CD4^+^ T_CM_ at day 13 post activation. **(D)** Cytokine staining comparing levels of IFN-γ, IL-17, IL-22 and IL-4 expression on *ex vivo* infected memory CD4^+^ T cells and non- polarized *in vitro* infection model T_CM_ at day 13 post activation. (**E)** TW10-specific CTL lamp-1 (CD107a) degranulation by flow cytometry on 2 donors after a 16h CTL challenge on uninfected cells (grey), HIV_JRSCF TW10esc_ -infected cells (blue) and HIV_JRCSF_-infected cells (red). (**F-G**) Sorting schematic to isolate HIV_JRSCF_TW10esc_ -infected and HIV_JRCSF_-infected infected cells after coculture (**F**) and without coculture (**G**) with TW10-specific CTL.

**Supplementary Figure 2**: **(A)** Abundance of recurring clonotypes from ECCITE-seq data of isolate HIV_JRSCF_TW10esc_ -infected and HIV_JRCSF_-infected infected cells with no CTL added, or as bystanders and survivors after a CTL killing assay from 1 donor. **(B)** Frequencies of mitochondrial genes in WNN clusters from infected survivor and bystander cell from all 4 donors. **(C)-(H)** Module scores of T-cell activation and functional state gene-sets from Szabo *et al*., 2019^36^ on bystander and survivor infected cells from all 4 donors.

**Supplementary Figure 3**: **(A)-(D)** Module scores of Hallmark OXPHOS, cholesterol homeostasis, fatty acid metabolism and hypoxia gene-sets on bystander and survivor infected cells from all 4 donors. P-values from Wilcoxon tests are sown.

**Supplementary Figure 4**: **(A)-(B)** Representative flow cytometry plots showing viability (top) and CD71 expression (bottom) on bulk CD4**^+^** T-cells after treatment with vehicle and different concentrations of DFO after 24 **(A)** or 72 **(B)** hours of treatment. **(C)** Representative and summary flow cytometry plots showing frequency of HIV-Gag expression in infected CD4**^+^** T-cells treated with vehicle or 10 µM DFO for 72 hours across 5 donors.

**Supplementary Figure 5**: **(A)-(B)** Violin plots showing expression of *BCL2* transcript and CD155 (PVR) surface protein expression on infected bystander and survivor cells across WNN clusters. **(C)** UMAP feature plot (left) and violin plot (right) showing surface protein expression of MHC-I (HLA-A-B-C) on infected bystander and survivor cells. **(D)** Violin plots showing surface protein expression of MHC-I on infected bystander and survivor cells across WNN clusters. **(E)** Ridge plots showing lower (MHC-I low), middle (MHC-I medium), and upper (MHC-I high) tercile surface protein expression of MHC-I across bystander and survivor cells. **(F)** UMAP projections showing cells clustered by WNN analysis and colored by bystander or survivor cell identity across MHC low, medium and high groups.

