## Supplementary figures and images for "Multi-Omic Atlas reveals cytotoxic phenotype and ROS-linked metabolic quiescence as key features of CTL-resistant HIV-infected CD4^+^ T-cells"

### Figure S1

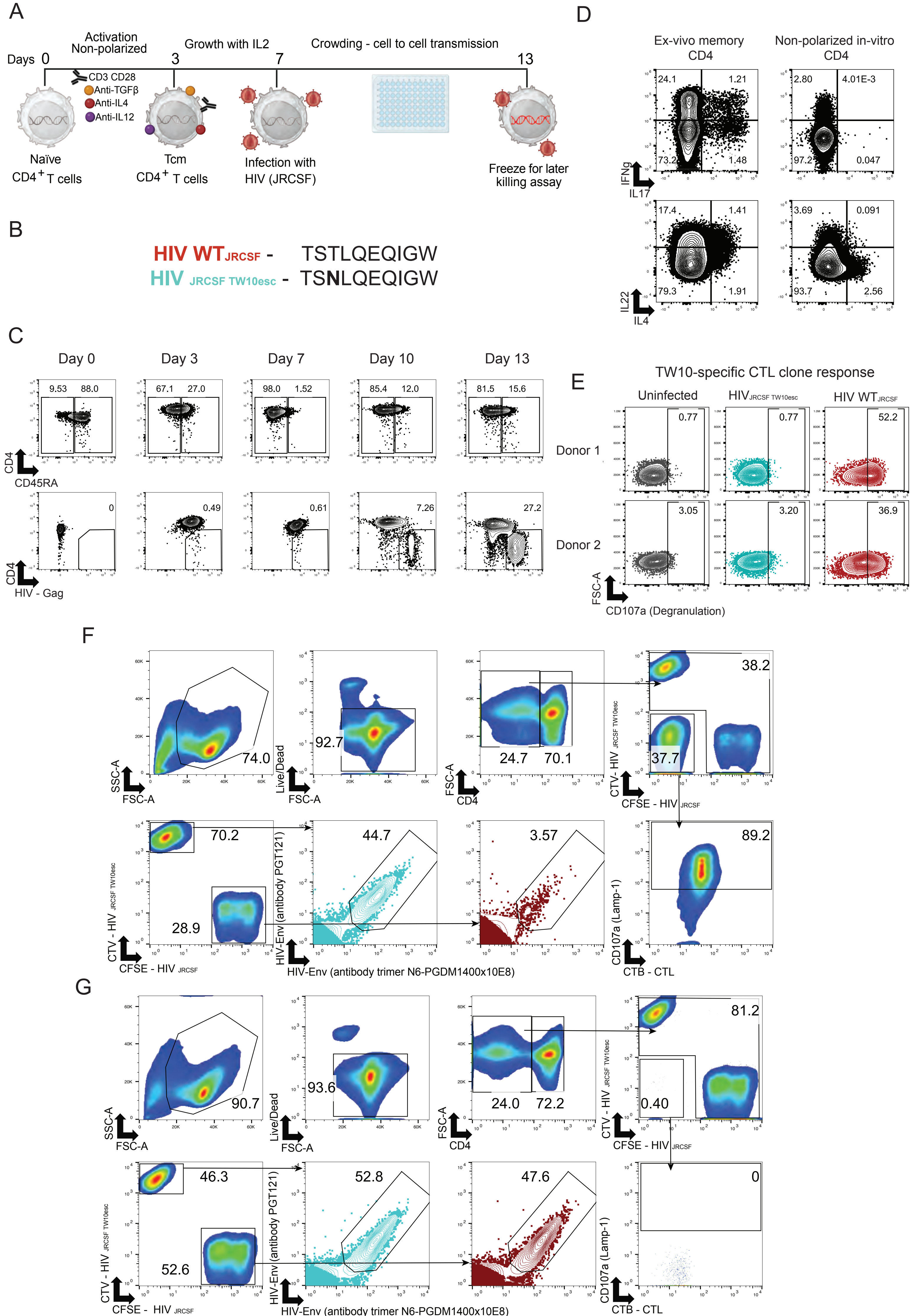

### Figure S2

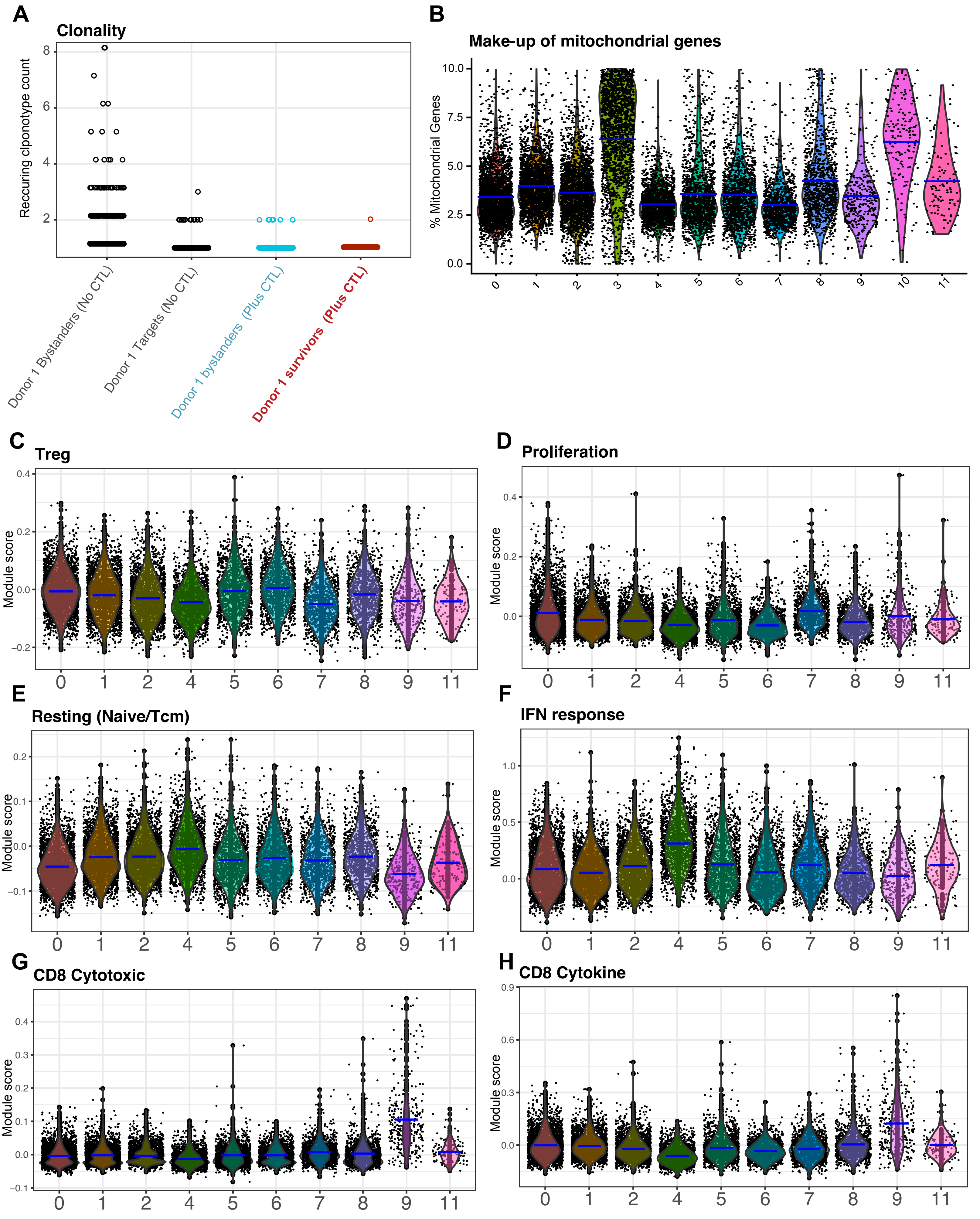

### Figure S3

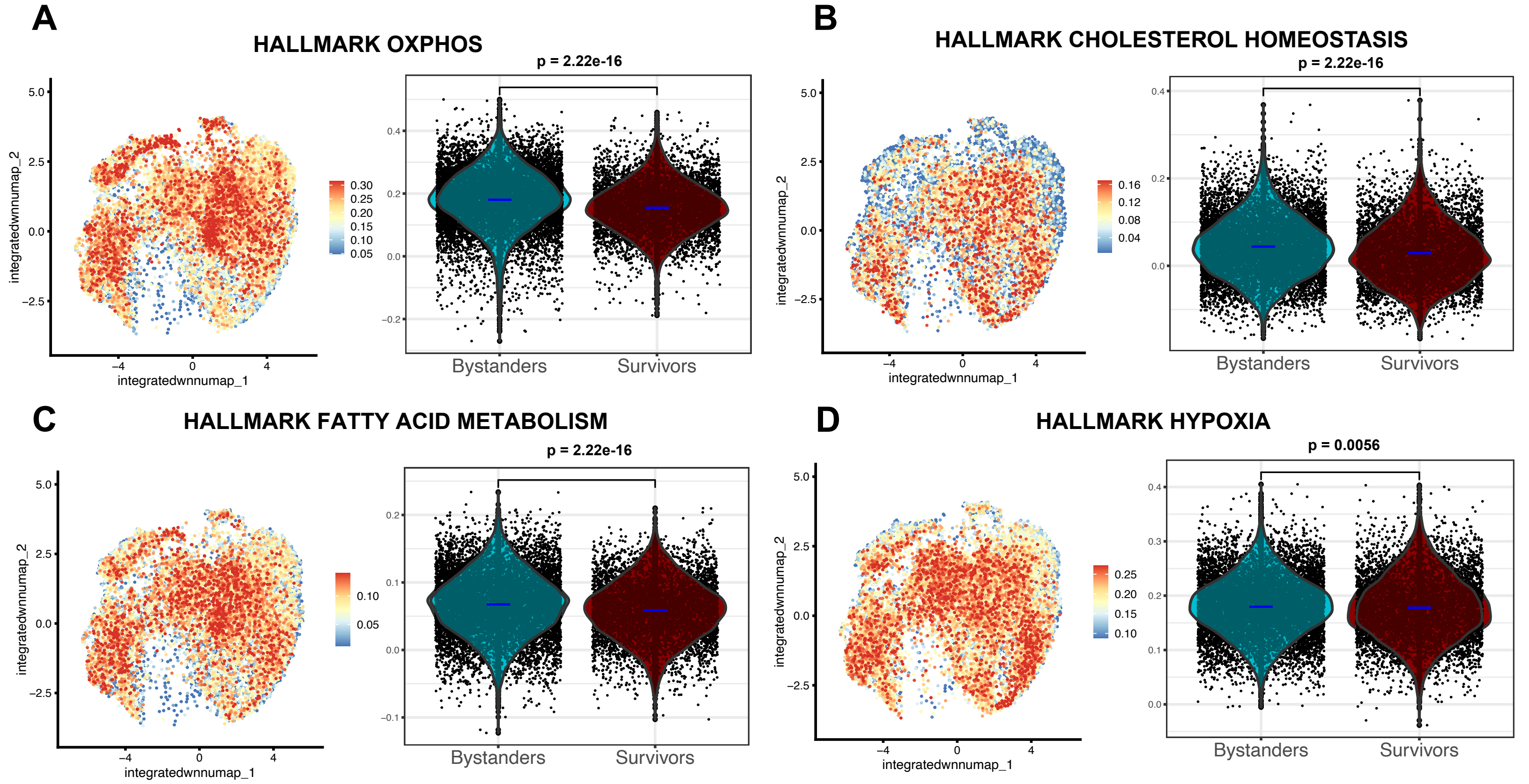

### Figure S4

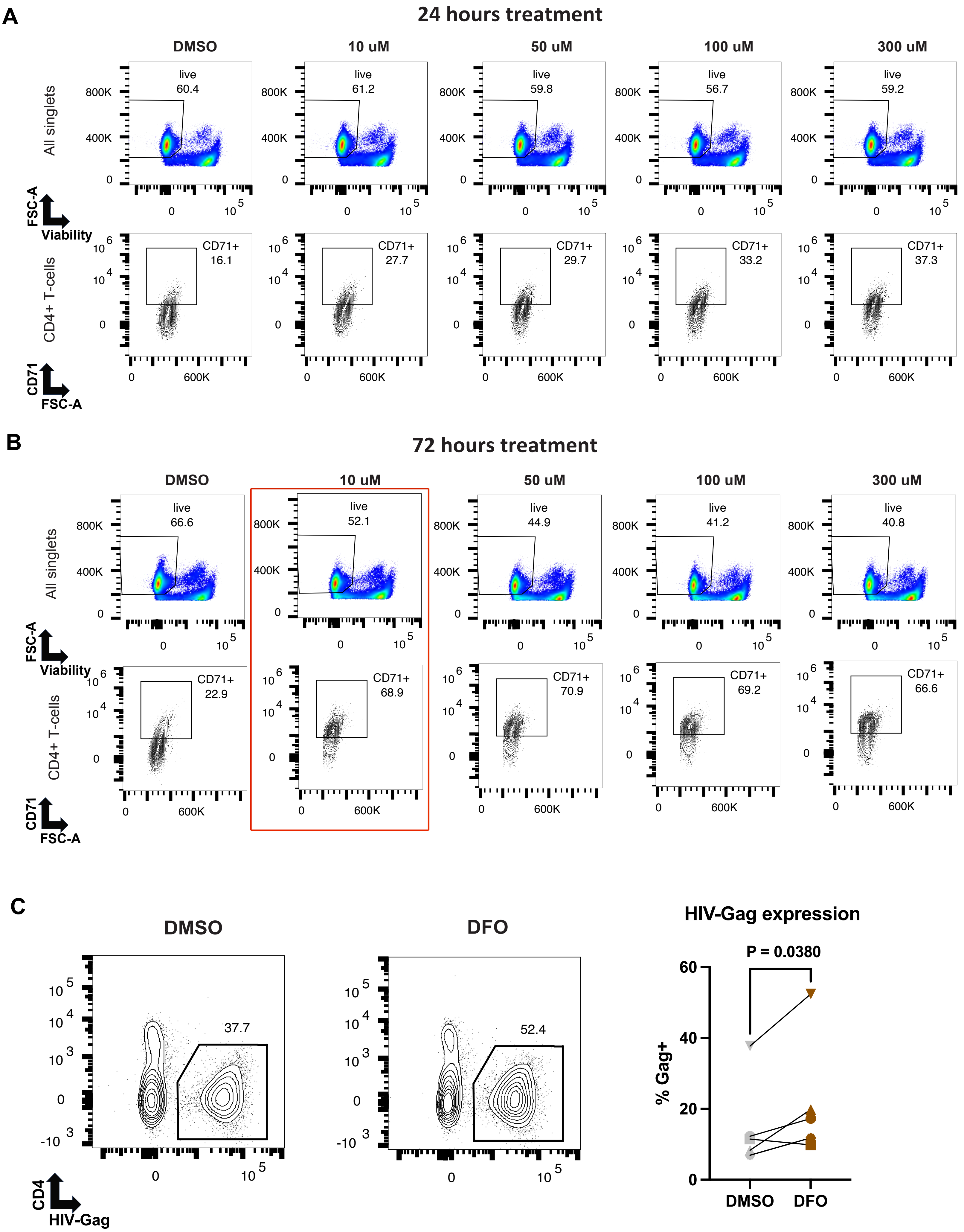

### Figure S5

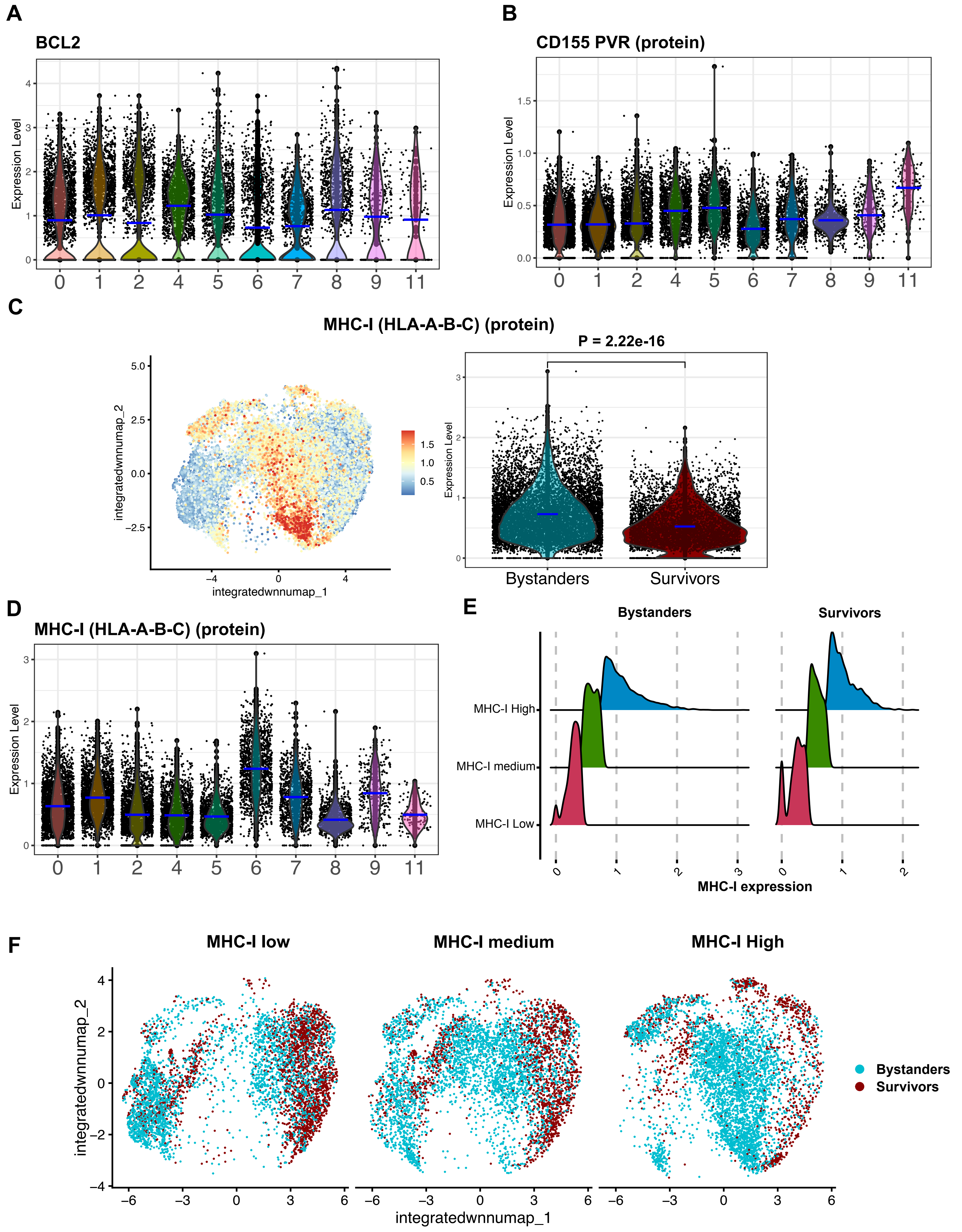
